# Convergent genomic variation and wild-type composite RNA-seq profiling identify candidate modules associated with 3-nitropropionic acid utilization in *Leclercia barmai*

**DOI:** 10.64898/2026.08.01.742209

**Authors:** Partha Barman, Shilpa Sinha, Subhajit Sen, Nibendu Mondal, Wriddhiman Ghosh, Ranadhir Chakraborty

## Abstract

3-Nitropropionic acid (3-NPA) is a toxic nitroaliphatic compound, but the genomic basis of bacterial adaptation to 3-NPA remains incompletely resolved, especially in organisms lacking a canonical nitronate monooxygenase. Here, we used phenotype screening, whole-genome resequencing, targeted UHPLC-MS/MRM, and composite RNA-seq profiling to investigate 3-NPA utilization in *Leclercia barmai* EMC7. Following attempted Tn5-Mob mutagenesis and kanamycin-based recovery, two independently derived mutants, LTM01 and LTM14, showed impaired growth under 3-NPA-dependent minimal conditions. No stable transposon insertion was detected in either mutant. Comparative genome analysis instead revealed convergent loss-of-function (LOF) mutations affecting amino acid metabolism, nitrogen allocation, central carbon metabolism, cofactor-linked functions, transport, and metal-redox homeostasis, while the predicted flavin-dependent nitro-redox candidate and canonical nitrate/nitrite reduction loci were not disrupted. Targeted UHPLC-MS/MRM analysis showed that wild-type EMC7 depleted approximately 50% of 3-NPA within 36 h, whereas the mutants displayed poor 3- NPA-utilization phenotypes under the same growth framework. Wild-type RNA-seq under glucose-3NPA relative to glucose-KNO3 identified a broad expression profile involving flavin-redox functions, nitrogen assimilation, oxidative-stress response, envelope stress, efflux, cofactor metabolism, and iron homeostasis. Integration of mutant LOF profiles with wild-type gene expression data showed that several disrupted genes or functional counterparts belonged to modules showing expression changes during the wild-type glucose-3NPA response. These findings identify candidate adaptive modules associated with 3-NPA utilization in EMC7, including nitrogen redistribution, carbon entry, sulfur-redox support, cofactor supply, transport, envelope remodeling, and metal-redox control. These results extend the landscape of metabolic adaptation beyond canonical pathways in environmental bacteria.

## 1. INTRODUCTION

3-Nitropropionic acid (3-NPA) is a naturally occurring nitroaliphatic compound produced by specific fungi and plants and is well known for its toxicity (Geddes et al., 2000; Hamilton et al., 2000; Reynolds and Lin, 2000; Upadhayay et al., 2023). In eukaryotic systems, 3-NPA acts as an irreversible inhibitor of succinate dehydrogenase, thereby disrupting respiration and energy metabolism (Francis et al., 2013; Reynolds and Lin, 2000; Scallet et al., 2003). In contrast, several microorganisms use 3-NPA as a growth substrate, often through oxidative nitroalkane metabolism involving nitronate monooxygenase-like or related flavin-dependent systems (Anderson et al., 2005; Barman et al., 2025; Becker et al., 2017; Francis et al., 2012; Nishino et al., 2010; Saha et al., 2019).

Genome annotation is widely used to connect microbial metabolic phenotypes with predicted enzymes. However, environmental bacteria often show substrate-utilization phenotypes that are not fully explained by canonical pathway annotation. Such cases indicate the involvement of non-canonical enzymes, distributed metabolic networks, or regulatory adaptations that link substrate transformation with nitrogen assimilation, carbon flux, stress tolerance, and redox balance (Huangyang and Simon, 2018; Snaebjornsson and Schulze, 2018).

*Leclercia barmai* EMC7, originally isolated from earthworm casting, is one such 3-NPA-utilizing environmental bacterium (Barman et al., 2025, 2022). Its genome encodes a flavin-dependent monooxygenase candidate, but the functional basis of 3-NPA utilization remains unresolved. Recent transcriptome-guided analysis of 3-NPA-degrading bacteria has shown that NPA metabolism is not always explained by canonical nitronate monooxygenase or pnoA-like expression alone. In *Pseudomonas* sp. Nvir, a flavin-dependent oxidoreductase was proposed as a candidate NPA-degrading enzyme despite weak support from canonical NMO-like candidates (Rogowska-van der Molen et al., 2022). These findings support a broader comparative microbial genomics view, in which flavin-redox functions, nitrogen-redox systems, and downstream assimilation modules are analyzed together during 3-NPA metabolism. This also makes EMC7 a useful system for examining whether 3-NPA utilization in environmental bacteria is associated with a discrete catabolic candidate or with broader metabolic and stress-response modules.

Forward genetic approaches provide a route to identify functional determinants of complex metabolic traits without prior assignment of a single candidate gene. Transposon mutagenesis is commonly used for this purpose because it is expected to generate insertional knockouts that link loss of phenotype to disruption of a specific locus (Bilyk et al., 2013; Cartman and Minton, 2010; Wall et al., 1996; Zhang et al., 2016). However, mutagenesis under strong selection can also reveal broader genome-level responses. Under strong selective pressure mutagenesis might not be restricted to discrete insertional events but can lead to genome-wide variation through stress-induced mutational processes (Foster, 2007, 2005; Frenoy and Bonhoeffer, 2018; Galhardo et al., 2007; Kivisaar, 2003; Matic, 2017; Wright, 2004). Such conditions may favour distributed mutations at multiple loci, especially in pathways involved in metabolism, stress response and cellular homeostasis. Thus, genotype– phenotype relationships that arise under these conditions may be due to perturbations at the network level rather than single gene effects.

Here, we used attempted Tn5-Mob mutagenesis as a phenotype-discovery approach to examine genomic variation associated with 3-NPA utilization in *L. barmai* EMC7. The expected outcome was recovery of insertional mutants in genes directly required for 3-NPA metabolism. Instead, two kanamycin-resistant, 3-NPA- impaired mutants lacked detectable stable transposon insertion and carried convergent chromosomal loss-of-function mutations in metabolic and stress-response modules. To resolve this non-canonical genotype-phenotype relationship, we combined phenotypic screening, targeted UHPLC-MS/MRM, whole-genome resequencing, and composite RNA-seq analysis of the wild-type bacteria. This comparative microbial genomics and transcriptome-supported framework was used to identify candidate modules associated with 3-NPA utilization in EMC7, including nitrogen allocation, carbon entry, sulfur-redox buffering, cofactor support, transport, and metal-redox homeostasis. Together, this study suggests that 3-NPA utilization in a non-model environmental bacterium is associated with multiple condition-responsive metabolic and stress-response modules, including nitrogen allocation, redox balance, and transport-associated functions.

## 2. MATERIALS AND METHODS

### 2.1 Bacterial strain

The 3-NPA-metabolizing strain EMC7, originally isolated from earthworm casting, was previously whole-genome sequenced using the Ion Torrent platform, assembled into contigs, annotated, and deposited in the NCBI database (Barman et al., 2025, 2022).

### 2.2 Kanamycin susceptibility testing of strain EMC7

The pSUP5011 suicide vector used for mutagenesis carries a kanamycin-resistance marker. Therefore, baseline kanamycin susceptibility of wild-type EMC7 was first confirmed to ensure that recovery on kanamycin-supplemented medium would not reflect intrinsic background resistance. The experiment was performed via the streak-plate method in Luria Agar (LA) plates, fortified with kanamycin (0.04 g/L). Plates were incubated at 30 °C, and the absence of visible growth was recorded after 48 h.

### 2.3 Electroporation-based delivery of pSUP5011:Tn5-Mob into EMC7 and phenotypic screening of electroporation-derived mutants

Random transposon mutagenesis of the kanamycin-sensitive strain EMC7 was attempted using the pSUP5011:Tn5-Mob system (Mukhopadhyaya et al., 2000; Simon et al., 1983). Following electroporation, kanamycin-containing medium was used for initial recovery of electroporation-derived mutants, while impaired 3-NPA utilization was identified by subsequent physiological screening on defined minimal media. Electrocompetent cells of strain EMC7 were prepared from an actively growing bacterial culture. Briefly, 0.1 mL of EMC7 seed culture was inoculated into 10 mL of LB broth and incubated at 30 °C with shaking at 100 rpm until the culture reached an OD_600_ of approximately 0.4, corresponding to ∼3.2 × 10^8^ cells/ mL. The culture was chilled on ice for 15–30 min and harvested by centrifugation at 2800 rpm for 15 min at 4 °C.

The cell pellet was first washed with 20 mL ice-cold, sterile double-distilled water and centrifuged at 2800 rpm for 20 min at 4 °C. The pellet was then washed with 10 mL ice-cold 10% glycerol, followed by four repeated washes with the same solution under identical centrifugation conditions. After the final wash, the cells were resuspended in 50 µL ice-cold 10% glycerol, centrifuged again, and finally resuspended in 50 µL ice-cold GYT medium (ESM_SI 1). A 1:10 dilution of the competent cell suspension was prepared in ice-cold GYT medium.

For electroporation, 40 µL of diluted competent EMC7 cells, corresponding to approximately 2.56 × 10 cells, was mixed with 2 µL of pSUP5011:Tn5-Mob plasmid DNA (50 ng/µL). The mixture was transferred to a pre-chilled 0.2 cm electroporation cuvette and pulsed once, 7.5 ms, using 2.5 kV, 300 Ω resistance, and 25 µF capacitance, corresponding to a field strength of 12.5 kV/cm (BIORAD MicroPulser electroporator).

Immediately after pulsing, the total content of the cuvette (∼40 µL) was transferred to a sterile tube containing 960 µL ice-cold SOC medium and kept at 30 °C for 1 h for post-electroporation recovery (ESM_SI 2). Following recovery, the cell suspension was further diluted by adding sterile 1.5 mL SOC at room temperature. An aliquot of 100 µL (cell suspension in SOC) per plate was spread onto each of the 25 SOB agar plates supplemented with kanamycin (0.04 g/L) (ESM_SI 3). Plates were incubated at 30 °C for 12 h.

A total of 1009 kanamycin-resistant CFUs were recovered. Details of the screening procedure have been given in the supplementary file (ESM_SI 4). Of this pool, two colonies that did not show growth on plates, composed of MSM supplemented with glucose, 3-NPA and kanamycin or MSM supplemented with 3-NPA but grew on plates composed of MSM supplemented with glucose, KNO_3_, and kanamycin or LA supplemented with kanamycin or LA, were clonally purified on MSM supplemented with glucose, KNO_3_, and kanamycin plates (ESM_SI 5; kanamycin used: 0.04 g/L) for further analysis.

### 2.4 Growth assay of wild-type EMC7 and the electroporation-derived mutants LTM01 and LTM14 in 3- NPA-supplemented MSM

Fresh single colonies of the two mutants and the wild-type bacterium were inoculated separately in MSM supplemented with glucose and KNO_3_ and grown overnight for 12 h (ESM_SI 5). The overnight grown cultures were inoculated (1% inoculum) separately into fresh MSM supplemented with 3- NPA or MSM supplemented with glucose and 3-NPA and incubated at 30 °C under shaking conditions (in an orbital shaker, 100 rpm) (ESM_SI 6 - SI 7). Cells from these cultures were withdrawn at regular intervals, suitably diluted in sterile MSM, and plated in Luria Agar (LA) plates for determining the viable cell counts. Finally, log CFU data was plotted against time. The study was performed in triplicate for statistical analysis.

Likewise, growth curves of the wild-type strain EMC7 and the two electroporation-derived mutants, LTM01 and LTM14, were derived from viable counts done at different time points over a period of 72 h in MSM supplemented with 3-NPA as the sole carbon and nitrogen source or 24 h in MSM supplemented with glucose (as sole carbon source) and 3-NPA (as sole nitrogen source).

### 2.5 Growth assay of wild-type EMC7 and the electroporation-derived mutants LTM01 and LTM14 in MSM supplemented with glucose and KNO_3_

Similar growth curve analysis of the mutants, LTM01 and LTM14, and the wild-type EMC7 was performed using viable counts at different time points over a period of 60 h in MSM supplemented with glucose (sole carbon source) and KNO_3_ (sole nitrogen source). The study was performed in triplicate for statistical analysis.

### 2.6 Heterotrophic growth study of the mutants and the wild-type in Luria broth

Growth curve analysis of mutants LTM01 and LTM14 and the wild-type EMC7 was performed using viable counts at different time points over a period of 6 h in Luria broth (M575-500G, HiMedia). The study was performed in triplicate for statistical analysis.

### 2.7 Targeted UHPLC-MS/MRM analysis of residual 3-NPA in EMC7 grown MSM supplemented with 3- NPA

Residual 3-NPA was quantified in wild-type EMC7 culture-supernatants (MSM supplemented with glucose and 3-NPA) using targeted UHPLC-MS analysis in multiple reaction monitoring (MRM) mode. The analysis was performed using an Acquity reverse-phase UHPLC system coupled with a Waters Xevo tandem quadrupole detector through an electrospray ionization source. Separation was performed on a Waters BEH C18 column, 50 x 2.1 mm, 1.7 µm. The mobile phase consisted of acetonitrile and buffer containing 5% methanol in water with 0.2% formic acid. The run used an isocratic 50:50 mobile phase composition for 3 min at a flow rate of 0.350 mL min^-1^. The column temperature was 35 ± 1 °C, sample temperature was 20 ± 2 °C, and injection volume was 2 µL.

3-NPA was monitored using the pure compound MRM signal at approximately 0.46 min with MRM transition m/z 118.0 46.0 in ESI-mode, with cone voltage of 25 V and collision energy of 8 V. The pure compound of 3- NPA was used for peak confirmation and calibration. Later, 3-NPA concentrations in EMC7 culture supernatants were calculated from the processed calibration curve. To visualize parent 3-NPA depletion, supernatants collected at 36 h were compared with 0 h 3-NPA reference sample, and reference-to-sample change was plotted. Abiotic controls were included to assess non-biological loss of 3-NPA, while *E. coli* K-12 (MTCC1302) was used as a non-3-NPA-utilizing biological control.

### 2.8 Determination of kanamycin MIC in wild-type EMC7 and electroporation-derived mutants LTM01 and LTM14

MIC of kanamycin was determined for LTM01 and LTM14 along with the wild-type strain EMC7 in Mueller Hinton Broth (HiMedia M391-500G) in the absence and presence of 0.1 g/L of Phenylalanine-arginine β-naphthylamide (PAβN) (Sen et al., 2022).

### 2.9 Growth response of EMC7, LTM01, and LTM14 under kanamycin exposure with and without PAβN

Finally, to evaluate the association of kanamycin-resistant phenotype of LTM01 and LTM14 with efflux-mediated tolerance under minimal-media conditions, growth of the two mutants (along with the wild-type) was monitored in MSM supplemented with glucose, KNO_3_, and kanamycin in the presence or absence of 0.1 g/L PAβN (ESM_SI 8). Overnight-grown cultures of EMC7, LTM01, and LTM14 were prepared separately in MSM supplemented with glucose and KNO_3_. These cultures were used as inocula (1%) into sterile MSM supplemented with glucose, KNO_3_, and kanamycin with or without PAβN. Cultures were incubated at 30 °C and 100 rpm for 72 h, and viable counts were determined on LA plates at 12 h intervals. Log CFU values were plotted against time. The experiment was performed in triplicate for statistical analysis.

### 2.10 Whole genome sequencing of electroporation-derived mutants LTM01 and LTM14

The mutants LTM01 and LTM14 were grown overnight in Luria broth at 30 °C and 100 rpm in an orbital shaker for genomic DNA isolation. Genomic DNA was extracted using the PureLink Genomic DNA Isolation Kit (Thermo Fisher Scientific, USA) and quantified with a SPECTROstar Nano microplate reader (BMG Labtech, Germany). Whole-genome shotgun sequencing was performed on an Ion S5 system (Thermo Fisher Scientific, USA), as described previously (Barman et al., 2022; Basak et al., 2021; Sen et al., 2020). High-quality reads were assembled into contigs via SPAdes v3.13.0 with default parameters (Bankevich et al., 2012). The raw sequencing data were deposited in the NCBI Sequence Read Archive. Draft genomes were annotated for open reading frames or coding sequences using NCBI Prokaryotic Genome Annotation Pipeline.

### 2.11 Screening for retained Tn5-Mob sequences

To assess whether LTM01 and LTM14 retained Tn5-Mob-derived sequences, both assembled mutant genomes were searched against the pSUP5011 sequence and relevant transposon-associated regions. The searched regions included Tn5-Mob-associated sequences, the kanamycin-resistance marker region, and adjacent vector-derived regions. Assembled contigs were screened using BLASTn-based sequence-similarity searches in BioEdit, with minimum identity and query coverage thresholds of 50% and 60%, respectively (Johnson et al., 2008). The low identity threshold was used as a permissive screen to avoid missing divergent or fragmented vector-derived sequences; candidate hits were retained only when supported by sufficient alignment coverage, fragment length, and read-level evidence. Associated raw sequencing reads were also mapped against the pSUP5011 reference sequence using BWA-MEM in Galaxy.eu platform to assess read-level support for retained Tn5-Mob-derived fragments. Candidate hits were evaluated based on alignment identity, query coverage, fragment length, and read support.

### 2.12 Genomic variant analysis in Galaxy.eu platform

Genomic variant analysis was performed on the Galaxy platform using the Snippy pipeline, followed by visualisation in JBrowse2 (Diesh et al., 2023). Raw single-end Ion-Torrent reads of the two mutant strains were quality-checked and aligned against the wild-type EMC7 reference genome supplied in FASTA format with its corresponding GFF/GBK annotation file. SNP and small indel calling were performed using Snippy with default parameters, using BWA-MEM for read mapping and FreeBayes for variant detection. Snippy-filtered variants were retained based on read depth, base quality, and allele frequency thresholds. The outputs included a core SNP alignment, VCF file, and GFF-formatted SNP annotation file describing variant positions and types (Connor et al., 2025).

As the Ion-Torrent sequencing is prone to small indel errors, particularly in homopolymeric regions, candidate insertion-deletion mutations were not accepted from automated calls alone. Biologically relevant variants were manually inspected in JBrowse2 using Snippy-generated BAM files, index files, and the SNP GFF file. The reference genome and annotation tracks were loaded first, followed by mapped-read and SNP tracks to assess coverage, alignment quality, and variant context. Variants were examined at gene-level resolution relative to coding sequences, promoter-proximal regions, and operon structures. Particular attention was given to mutations affecting antibiotic resistance, envelope stress response, carbon and nitrogen metabolism, porin-mediated permeability, transcriptional regulation, and efflux transport. Variant coordinates and gene-level impacts were manually curated for downstream genotype-phenotype interpretation and figure preparation. Promoter prediction of the ompC upstream sequence in LTM01 and EMC7 was performed using the BDGP Neural Network Promoter Prediction server in prokaryotic mode with a 0.8 cut-off value (Reese, 2001).

### 2.13 Composite RNA-seq analysis of wild-type EMC7 grown under glucose-3NPA and glucose-KNO_3_ conditions

Composite RNA-seq analysis was performed to profile the transcriptional response of wild-type (WT) *Leclercia barmai* strain EMC7 under 3-nitropropionic acid exposure. WT EMC7 was grown to mid log phase in minimal salt medium supplemented with glucose and KNO_3_ (glucose-KNO_3_) as the control condition or in minimal salt medium supplemented with glucose and 3-nitropropionic acid (glucose-3NPA) as the test condition. The comparison was aimed to identify transcriptional modules associated with 3-NPA response in a common glucose background.

The experiment was performed using three independent cultures for each condition. Equal volumes of cultures from the three replicates were pooled before RNA extraction to prepare one composite RNA sample for the glucose-KNO_3_ control condition and one composite RNA sample for the glucose-3NPA test condition. Total RNA was extracted from the composite samples and subjected to quality assessment before library preparation. RNA integrity was evaluated using an Agilent TapeStation system with RNA ScreenTape. RNA samples that passed quality control were processed for library preparation. Composite RNA samples were used to capture condition-level transcriptional trends, but gene-level statistical inference should be interpreted with caution. RNA libraries were prepared after RNA fragmentation, followed by library quality assessment using Agilent D1000 ScreenTape. Libraries were then sequenced on an Illumina NovaSeq 6000 platform using 150 bp paired-end sequencing chemistry.

Raw sequencing reads were subjected to initial quality control using FastQC, and quality reports were summarized using MultiQC. Adapter sequences and low-quality bases were removed using Trim Galore. Read trimming was performed with a Phred score threshold greater than 20 and a minimum retained read length of 20 bp. The processed reads were aligned to the EMC7 reference genome using HISAT2. Gene-level read counts were generated for downstream comparative expression profile analysis.

Expression profile comparison was conducted using iGEAK, with glucose-3NPA treated as the test condition and glucose-KNO_3_ treated as the control condition. Genes showing expression differences with an applied fold-change threshold of > 1.5 and p ≤ 0.05 were retained for functional interpretation. The differentially expressed genes were functionally grouped by annotation into modules associated with flavin-dependent oxidoreduction, nitro-group redox handling, nitrate/nitrite sensing, nitrate reductase-associated functions, molybdenum cofactor metabolism, nitrogen regulation, redox and respiratory metabolism, oxidative stress mitigation, Fe-S cluster maintenance, DNA repair, envelope stress response, efflux, porin regulation, and central carbon metabolism.

The WT transcriptome was used as a response map to interpret the genomic lesions identified in the 3-NPA- impaired mutants LTM01 and LTM14. We compared WT glucose-3NPA-responsive genes showing expression differences with common loss-of-function mutations identified in both mutants. Since transcriptome identifiers and variant-derived gene identifiers were not always directly matched at single-locus level, overlap was interpreted at the annotation and functional-module level, rather than as confirmed one-to-one gene identity. This comparison was used to assess whether shared mutant lesions mapped to modules showing expression changes in WT EMC7 during glucose-3NPA exposure.

### 2.14 Statistical analysis

Statistical analyses were performed using ANOVA followed by Tukey’s HSD test in IBM_SPSS v29.0 suite (Field, 2024).

## 3. RESULTS

### 3.1 Tn5-Mob-based screening recovered electroporation-derived mutants with impaired 3-NPA- dependent growth

Using Tn5-Mob mediated mutagenesis followed by kanamycin selection, colonies were screened based on their ability to grow in minimal medium with 3-NPA as the sole nitrogen source. Two independent electroporation-derived mutants, designated LTM01 and LTM14, consistently exhibited impaired growth under these conditions. These mutants, however, were able to grow on media enriched with nitrate (glucose-KNO_3_-MSM) and on complex nutrient rich medium (LB) (Fig. 1a-1d).

**Fig. 1.**
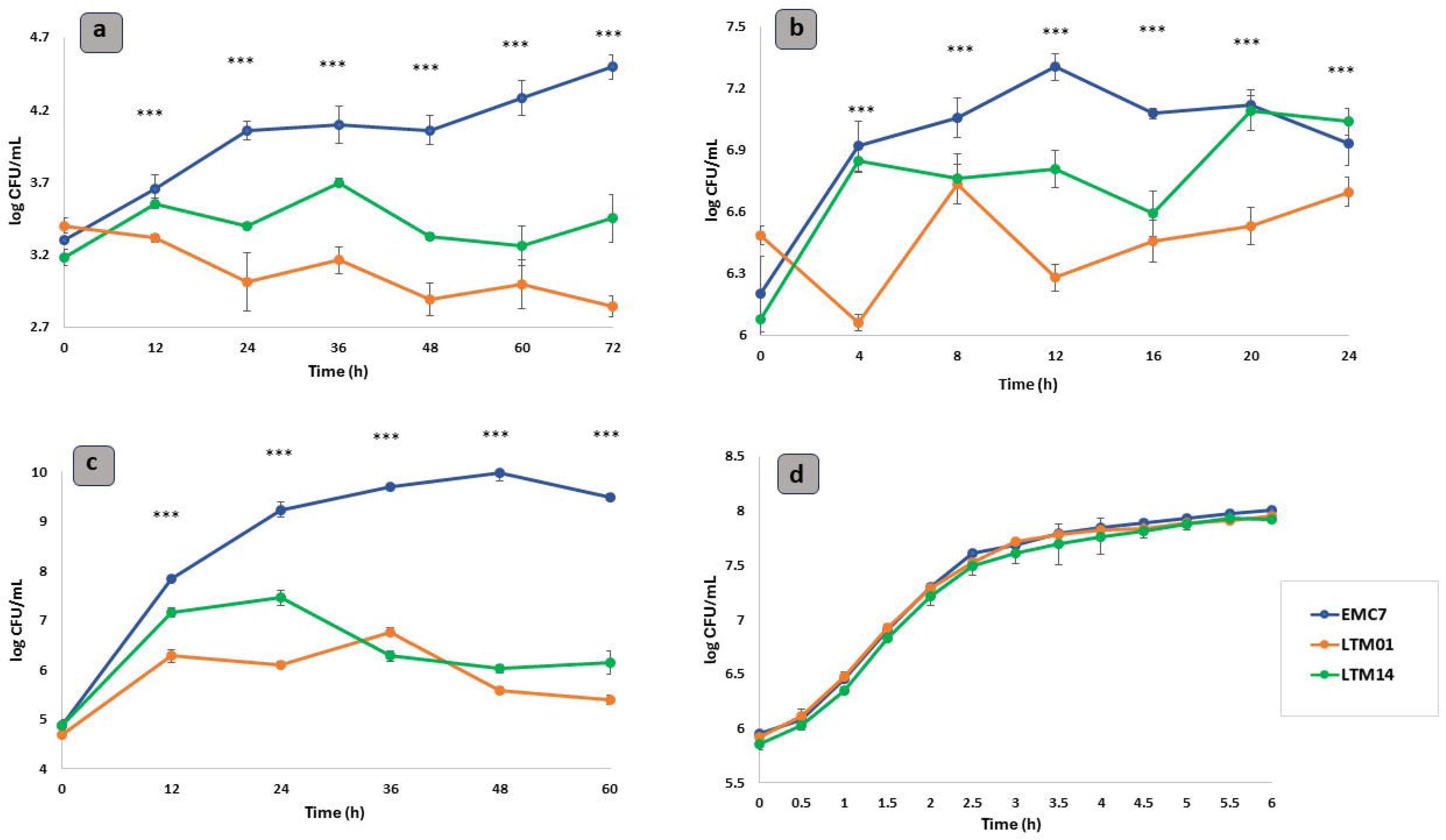
Phenotypic characterization in terms of growth response of EMC7, LTM01, and LTM14 under varying carbon and nitrogen sources in minimal salt medium (n = 3). a) MSM supplemented with 3-NPA as sole carbon and nitrogen source. Growth differed significantly among strains, time points, and strain × time interaction by two-way ANOVA. Tukey HSD separated all three strains, with EMC7 showing the highest growth, LTM14 intermediate growth, and LTM01 the lowest growth. Pairwise strain comparisons were significant for EMC7 vs LTM01, mean difference = 0.8816, p < 0.001; EMC7 vs LTM14, mean difference = 0.5382, p < 0.001; and LTM14 vs LTM01, mean difference = 0.3433, p < 0.001. *** indicates p < 0.001 among strain-level comparisons in this condition. b) MSM supplemented with Glucose (as sole carbon source) and 3-NPA (as sole nitrogen source). Two-way ANOVA showed significant effects of strain, time, and strain × time interaction. EMC7 showed the highest overall growth, followed by LTM14 and LTM01. Tukey HSD showed significant differences for EMC7 vs LTM01, mean difference = 0.4876, p < 0.001; EMC7 vs LTM14, mean difference = 0.2418, p < 0.001; and LTM14 vs LTM01, mean difference = 0.2459, p < 0.001. *** indicates p < 0.001 among strain-level comparisons in this condition. c) MSM supplemented with Glucose (as sole carbon source) and KNO_3_ (as sole nitrogen source). Growth differed strongly among the three strains. Two-way ANOVA showed significant effects of strain, time, and strain × time interaction. Tukey HSD showed that EMC7 had significantly higher growth than both mutants, while LTM14 was significantly higher than LTM01. Pairwise comparisons were EMC7 vs LTM01, mean difference = 2.641, p < 0.001; EMC7 vs LTM14, mean difference = 2.164, p < 0.001; and LTM14 vs LTM01, mean difference = 0.477, p < 0.001. *** indicates p < 0.001 among strain-level comparisons in this condition. d) Heterotrophic medium (Luria broth). All strains showed rapid growth and reached a near-plateau by 4 to 6 h. Two-way ANOVA showed significant strain and time effects, but the strain × time interaction was not significant, F(24,78) = 0.363, p = 0.997. Post hoc comparison showed no significant difference between EMC7 and LTM01, mean difference = 0.020, p = 0.782. LTM14 was significantly lower than EMC7, mean difference = 0.084, p < 0.001, and lower than LTM01, mean difference = 0.063, p = 0.002. Thus, no time-specific divergence is indicated in LB medium.

Both mutants showed reduced cell density cultured in minimal medium containing glucose and 3-NPA when compared to the wild-type strain EMC7. This defect was more pronounced when 3-NPA was used as both carbon and nitrogen source (Fig. 1a, 1b). In Luria broth, all strains showed rapid growth and reached a similar near-plateau by 4 to 6 h, with no time-specific divergence among strains (Fig. 1d).

Growth of EMC7, LTM01, and LTM14 was also evaluated in minimal salt medium containing glucose with either nitrate or 3-NPA as the nitrogen source to evaluate nitrogen utilization.

In glucose-KNO_3_ medium (when evaluated for nitrogen utilization), all three strains were able to grow, but the mutants showed altered growth kinetics relative to wild-type EMC7, with LTM01 showing the stronger reduction and LTM14 showing an intermediate profile (Fig. 1c). In the 3-NPA medium, both mutants showed consistently lower growth than wild-type EMC7 across the observation period (Fig. 1b). However, when 3-NPA was utilized as the sole source of both carbon and nitrogen, the growth inhibition in the mutants was significantly more pronounced (Fig. 1a).

### 3.2 Targeted UHPLC-MS/MRM showed 3-NPA depletion by WT EMC7

Residual 3-NPA in culture supernatants was analyzed using targeted UHPLC–MS/MRM. The starting concentration of 3-NPA was 59.95 mg L ¹ at 0 h. After 36 h of incubation, the concentration in cultures with wild-type EMC7 decreased to 30.22 mg L ¹. This corresponded to approximately 49.6% depletion of 3-NPA. In contrast, abiotic controls, as well as *E. coli* K-12 cultures, demonstrated no significant changes in 3-NPA concentration during the same experimental timeframe (shown in ESM_Supplementary Fig. S2a-2b).

### 3.3 Kanamycin resistance in LTM01 and LTM14 was associated with PA**β**N-sensitive tolerance

Minimum inhibitory concentration (MIC) assays showed that LTM01 and LTM14 had higher kanamycin resistance than wild-type EMC7 (Fig. 2). Wild-type EMC7 showed a kanamycin MIC of 10 µg mL ¹, whereas LTM01 and LTM14 showed MIC values of 80 and 90 µg mL ¹, respectively. In the presence of PAβN, the MIC of wild-type EMC7 remained unchanged at 10 µg mL ¹, while the MIC values of LTM01 and LTM14 decreased to 70 and 70 µg mL ¹, respectively. This PAβN-associated reduction supports an efflux-linked contribution to kanamycin tolerance in the mutants. Growth analysis under kanamycin exposure with or without PAβN further showed altered mutant growth kinetics relative to wild-type EMC7 (ESM_Supplementary Fig. S1). These results supported kanamycin resistance as a secondary selected phenotype that required genomic explanation.

**Fig. 2.**
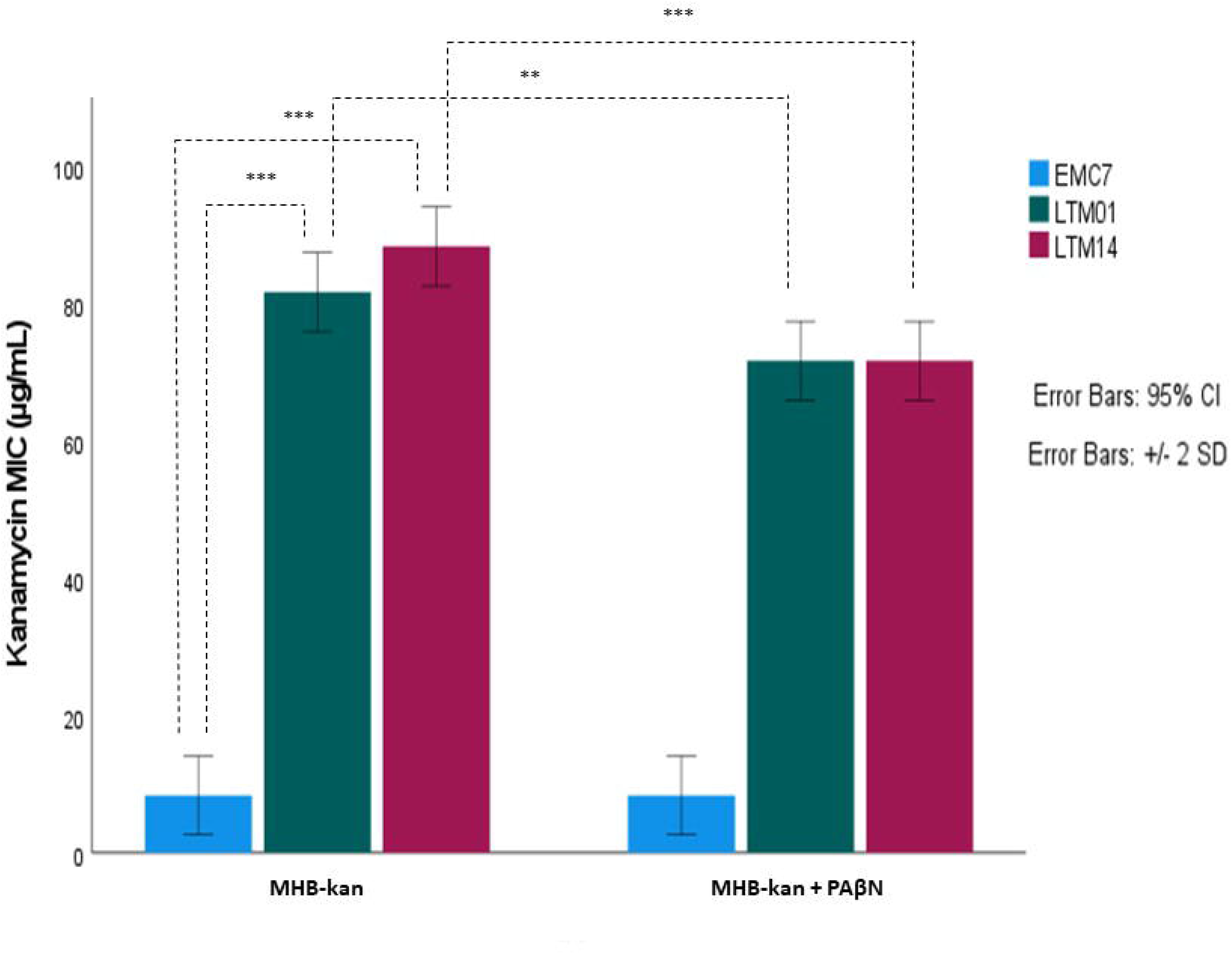
Minimum inhibitory concentrations of kanamycin for strains EMC7, LTM01, and LTM14 in the presence and absence of PAβN in MHB medium (n = 3). Bars represent mean MIC values from three independent replicates, and error bars showing 95% confidence intervals. Two- way ANOVA showed significant effects of strain, condition, and strain × condition interaction on kanamycin MIC. PAβN treatment reduced the MIC of LTM01 from 80 to 70 µg mL ¹ and LTM14 from 90 to 70 µg mL ¹, while EMC7 remained unchanged at 10 µg mL ¹. This pattern indicates a partial efflux-associated contribution to kanamycin tolerance in the mutants. Asterisks indicate significant pairwise differences, **p < 0.01, ***p < 0.001.

### 3.4 Electroporation-derived mutants lacked detectable stable Tn5-Mob insertion

Whole-genome sequencing was performed to identify genomic changes associated with the stable 3-NPA- impaired and kanamycin-resistant phenotypes of LTM01 and LTM14. No full-length Tn5-Mob insertion, junction-supported insertion site, or reliably supported partial Tn5-Mob-derived fragment was detected in either assembled contigs or raw sequencing reads from LTM01 and LTM14.

### 3.5 Shared loss-of-function (LOF) mutations occurred in genes linked to nitrogen, carbon-redox, cofactor, transport, and metal-homeostasis modules

Upon detailed sequence inspection, multiple insertion–deletion (indel) mutations were identified in both mutant genomes. Several loss-of-function mutations were shared between LTM01 and LTM14, including genes associated with amino acid metabolism and transport, such as *asnB, gcvA, sdaA, sdaB, cysE, ilvY, hisM,* and *glnQ* (Table 2).

**Table 1.**
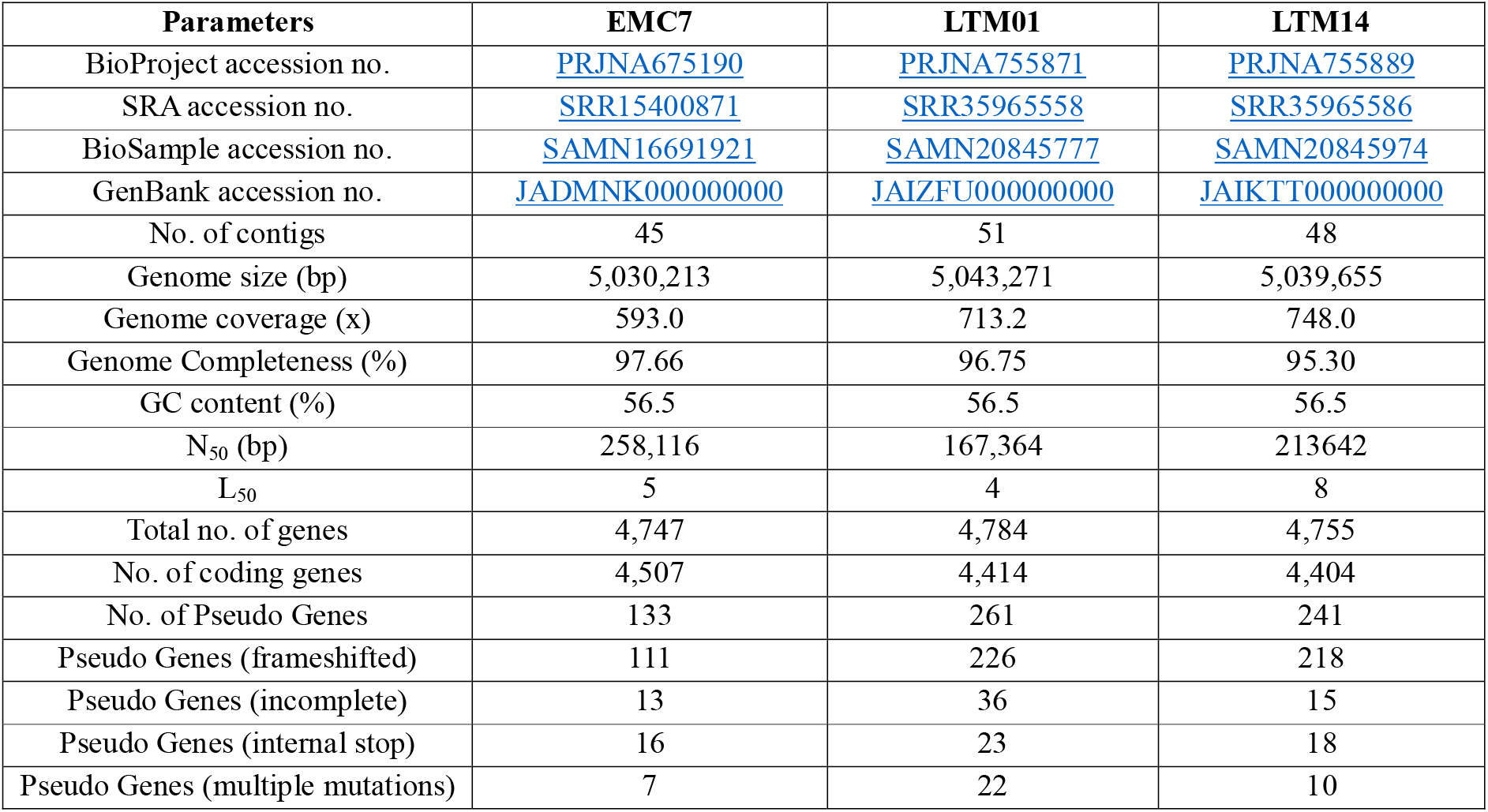
Genome assembly and annotation statistics of *Leclercia barmai* strain EMC7 and mutants LTM01 and LTM14, sequenced using the Ion S5 system and assembled using SPAdes3.13.0.

| Parameters | EMC7 | LTM01 | LTM14 |
| --- | --- | --- | --- |
| BioProject accession no. | PRJNA675190 | PRJNA755871 | PRJNA755889 |
| SRA accession no. | SRR15400871 | SRR35965558 | SRR35965586 |
| BioSample accession no. | SAMN16691921 | SAMN20845777 | SAMN20845974 |
| GenBank accession no. | JADMNK000000000 | JAIZFU000000000 | JAIKTT000000000 |
| No. of contigs | 45 | 51 | 48 |
| Genome size (bp) | 5,030,213 | 5,043,271 | 5,039,655 |
| Genome coverage (x) | 593.0 | 713.2 | 748.0 |
| Genome Completeness (%) | 97.66 | 96.75 | 95.30 |
| GC content (%) | 56.5 | 56.5 | 56.5 |
| N<sub>50</sub> (bp) | 258,116 | 167,364 | 213,642 |
| L<sub>50</sub> | 5 | 4 | 8 |
| Total no. of genes | 4,747 | 4,784 | 4,755 |
| No. of coding genes | 4,507 | 4,414 | 4,404 |
| No. of Pseudo Genes | 133 | 261 | 241 |
| Pseudo Genes (frameshifted) | 111 | 226 | 218 |
| Pseudo Genes (incomplete) | 13 | 36 | 15 |
| Pseudo Genes (internal stop) | 16 | 23 | 18 |
| Pseudo Genes (multiple mutations) | 7 | 22 | 10 |

**Table 2.**
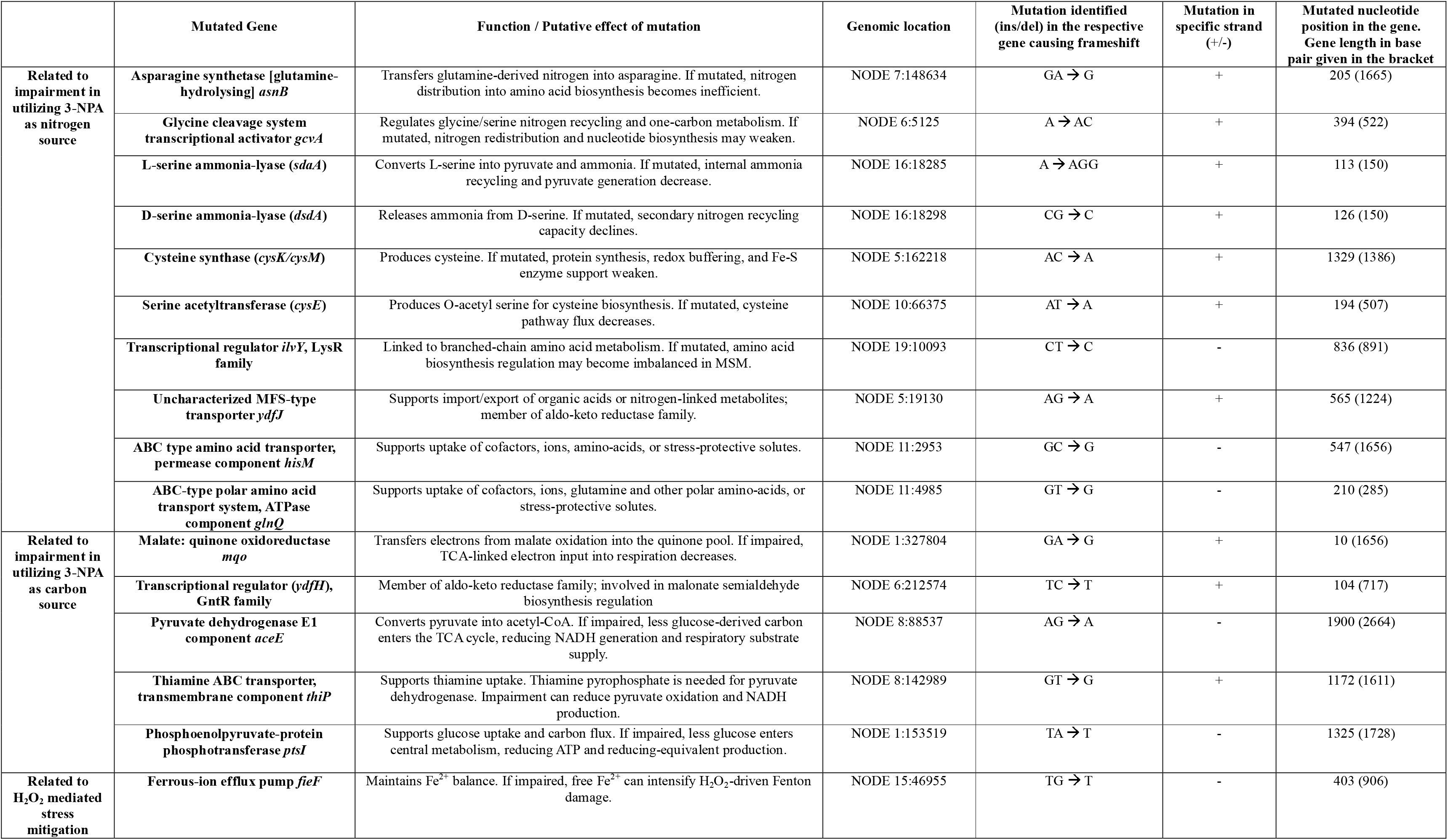
Shared loss-of-function mutations linked to impaired 3-NPA-dependent growth in mutant strains. Mutated genes were grouped by their predicted roles in nitrogen assimilation, carbon flux, and oxidative stress control.

More mutations affecting genes linked to central metabolism and cofactor-related processes, including *aceE, ptsI,* and *thiP*, were also detected. Mutations in genes associated with metal ion transport, including *fieF*, were observed in both mutants (Table 2).

LTM01 contained additional frameshift mutations affecting oxidoreductases and regulatory proteins, including *yagR*, *nasA*, and multiple transcriptional regulators (ESM_Supplementary Fig. S3a–S3g, S4a, S4b). LTM14 showed fewer such mutations, with distinct alterations in regulatory regions (ESM_Supplementary Fig. S5, S6a, S6b).

### 3.6 WT EMC7 transcriptome identified a broad glucose-3NPA-associated expression profile

RNA-seq analysis was performed to compare gene expression in wild-type EMC7 grown in minimal medium containing glucose with either 3-NPA or nitrate as the nitrogen source.

Sequencing generated 35,180,234 raw reads for the glucose-KNO_3_ control sample and 35,092,982 raw reads for the glucose-3NPA sample. After trimming, 35,179,740 and 35,092,146 reads were retained for glucose-KNO_3_ and glucose-3NPA, respectively. Alignment of trimmed reads to the EMC7 reference genome showed high mapping rates of 94.72% for glucose-KNO_3_ and 97.97% for glucose-3NPA. Properly paired reads accounted for 89.83% and 94.65% of reads in the glucose-KNO_3_ and glucose-3NPA samples, respectively.

A total of 1,279 genes showed expression differences under glucose-3NPA relative to glucose-KNO_3_, including 492 upregulated and 787 downregulated genes (ESM_Supplementary Fig. S7a-S7c). Upregulated genes included those encoding flavin-dependent oxidoreductases, nitroreductases, monooxygenase-like proteins, and components of nitrate/nitrite sensing and reduction systems.

Genes involved in redox processes, including NADH dehydrogenase, catalase, glutathione transferase, and thioredoxin-related proteins, were also upregulated. In addition, genes associated with envelope stress response and efflux systems, including RND transporters and TolC, showed increased expression (ESM_SI9).

Downregulated genes included those encoding major outer membrane porins (ompC, ompA), lipoproteins, and components associated with membrane structure and permeability. Several genes related to carbohydrate metabolism and specific respiratory pathways were also downregulated (ESM_SI 10–11).

### 3.7 Shared LOF mutations mapped to WT glucose-3NPA-responsive modules in *L. barmai*

Comparison of loss-of-function mutations in LTM01 and LTM14 with genes showing expression differences in wild-type EMC7 under glucose-3NPA conditions identified corresponding functional categories. Several genes mutated in both mutants corresponded to genes upregulated in the wild-type transcriptome during growth on 3- NPA. These included genes associated with amino acid metabolism, nitrogen regulation, and metabolite transport (Table 3).

**Table 3.** Module-level correspondence between shared LOF mutations and WT EMC7 glucose-3NPA- associated expression profiles (with associated adjusted p values ranging from 2.92 × 10^²³^ to 1.35 × 10)

| Shared LOF gene in mutants | Relevance | Functional module | Gene ID in wild-type transcriptome | Log2FC |
| --- | --- | --- | --- | --- |
| *asnB* | 3NPA-derived nitrogen handling | Nitrogen allocation | asparagine synthetase B, ITX56_14090 | +2.188 |
| *gcvA* | 3NPA-derived nitrogen handling | Glycine and one-carbon regulation | glycine cleavage system transcriptional regulator GcvA, ITX56_12310 | +2.405 |
| *sdaA* | 3NPA-derived nitrogen handling | L-serine nitrogen and pyruvate generation | L-serine ammonia-lyase, ITX56_12330; L-serine ammonia-lyase, ITX56_03275 | +2.595; +3.136 |
| *sdaB* | 3NPA-derived nitrogen handling | D-serine nitrogen recycling | D-serine ammonia-lyase, ITX56_21405 | +2.535 |
| *cysE* | 3NPA-derived nitrogen handling | Cysteine pathway entry | serine acetyltransferase, ITX56_18175 | +3.066 |
| *ilvY* | 3NPA-derived nitrogen handling | Regulatory remodeling | HTH-type transcriptional activator IlvY, ITX56_22655 | +2.452 |
| *ydfJ* | 3NPA-derived nitrogen handling | MFS-transporters | ITX56_00445, ITX56_05945, ITX56_08095 | +1.67 to +3.54 |
| *hisM, glnQ* | 3NPA-derived nitrogen handling | Amino acid transport | amino acid ABC/permease transporters | +1.56; +3.91 |
| *aceE* | 3NPA-derived carbon processing | Pyruvate to acetyl-CoA entry | pyruvate dehydrogenase, ITX56_14955; PDH E2 component, ITX56_14950 | +2.088; +2.154 |
| *thiP* | 3NPA-derived carbon processing | Thiamine-linked cofactor support | thiamine/thiamine pyrophosphate ABC transporter permease ThiP, ITX56_15195 | +2.12 |
| *ptsI* | 3NPA-derived carbon processing | PTS-mediated carbon entry | phosphoenolpyruvate-protein phosphotransferase PtsI, ITX56_00685 | 2.062 |
| *fieF* | ROS mitigation | Iron and metal-redox control | CDF family cation-efflux transporter FieF, ITX56_20995 | 2.097 |

Similarly, genes involved in central carbon metabolism and cofactor-dependent processes, including *aceE*, *ptsI*, and *thiP*, were both mutated in the mutants and upregulated in the wild-type under 3-NPA conditions. Mutations in the metal ion transport gene *fieF* were also observed in both mutants, corresponding to genes showing altered expression in the wild-type transcriptome.

## 4. DISCUSSION

This study examined 3-NPA utilization in the environmental bacterium *Leclercia barmai* EMC7 by combining mutant phenotyping, targeted UHPLC-MS/MRM, whole-genome resequencing, screening for retained Tn5-Mob sequences, and wild-type composite RNA-seq profiling. The central observation was the recovery of kanamycin-resistant, 3-NPA-impaired electroporation-derived mutants in the absence of detectable stable Tn5- Mob insertion. Instead of identifying a single disrupted catabolic locus, the combined genomic and transcriptomic evidence identified candidate modules associated with nitrogen allocation, carbon entry, cofactor support, transport, envelope remodeling, and redox homeostasis (Fig. 3). Thus, the present dataset supports a candidate network-level model for 3-NPA utilization in EMC7, while recognizing that individual causal relationships require direct functional validation.

**Fig. 3.**
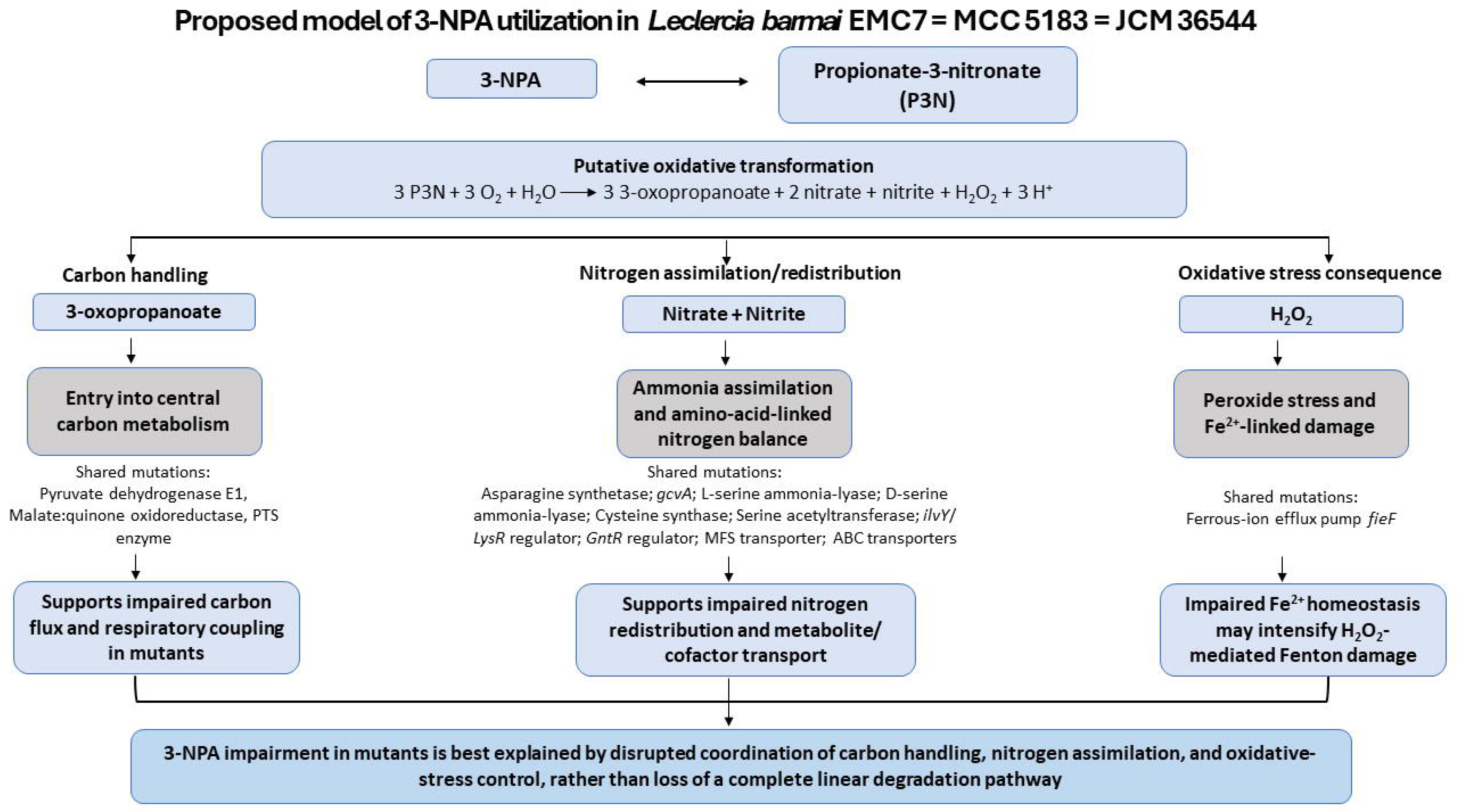
Proposed working model for candidate modules associated with 3-NPA utilization in *Leclercia barmai* EMC7. The model summarizes phenotypic impairment in LTM01 and LTM14, convergent genomic variation, and wild-type composite RNA-seq-supported modules. The model is hypothesis-driven and indicates candidate links among carbon handling, nitrogen redistribution, oxidative-stress response, transport, and metal-redox homeostasis.

The absence of full-length, junction-supported, or reliably supported partial Tn5-Mob-derived sequences in both assembled contigs and raw reads argues against retained canonical Tn5-Mob insertional knockouts as the basis of the stable 3-NPA-impaired and kanamycin-resistant phenotypes of LTM01 and LTM14. This outcome is consistent with a non-canonical mutagenesis-associated process in which transient transposase activity, DNA damage, selection-associated repair, or selection of pre-existing variants contributed to stable chromosomal variation rather than retained transposon insertion. Although highly fragmented or rearranged transposon-derived sequences cannot be completely excluded, the recovery of similar phenotypes in independently derived mutants, together with genome-wide sequence analysis, supports the interpretation that the observed phenotypes are associated with stable chromosomal changes rather than canonical insertional knockouts. This interpretation is consistent with previous reports of abortive or “hit-and-run” transposition events (Bhasin et al., 1999; Jilk et al., 1993; Li et al., 2020; Reznikoff, 2003) and highlights the need to interpret mutagenesis-derived phenotypes in non-model bacteria beyond simple single-gene disruption models.

A key finding of this work is that impaired 3-NPA utilization in LTM01 and LTM14 was not linked to mutation of the predicted flavin-dependent nitro-redox candidate gene (MBZ0056881.1) or canonical assimilatory nitrate/nitrite reduction loci. This is not consistent with a simple proximal block in substrate conversion. Instead, shared loss-of-function mutations affected genes involved in amino acid metabolism, nitrogen redistribution, cofactor-linked metabolism, transport, and metal-redox homeostasis. These patterns suggest that the poor 3-NPA growth phenotype is associated with disruption of support modules potentially involved in integrating 3-NPA- derived nitrogen and downstream metabolic stress into cellular physiology. Altered mutant growth kinetics under nitrate-dependent minimal conditions further suggest that the altered phenotype may not be restricted to 3- NPA exposure alone, but extends to broader nitrogen allocation under nutrient-limited conditions.

The wild-type composite RNA-seq profile further supports this module-level interpretation. Growth under glucose-3NPA relative to glucose-KNO3 was associated with broad transcriptional remodeling involving flavin-dependent redox enzymes, nitrate/nitrite-responsive systems, respiratory components, oxidative-stress defence, envelope stress, efflux, and membrane-remodeling functions. The induction of flavin monooxygenase-like systems and related redox enzymes suggests possible involvement of non-canonical nitroaliphatic transformation pathways, although direct enzymatic roles remain to be established (Rogowska-van der Molen et al., 2022). Because the RNA-seq analysis was based on composite samples, these data are best interpreted as a condition-level transcriptional response profile rather than replicate-resolved gene-level proof. Within that scope, the expression profile indicates that glucose-3NPA growth is associated with nitrogen-redox and cofactor-associated functions distinct from the glucose-KNO_3_ control condition.

A notable feature of the 3-NPA response was the coordinated expression of tolerance-associated functions such as efflux systems and reduced membrane permeability. The WT transcriptome showed increased expression of RND efflux components and TolC, together with reduced expression of major porins, suggesting that EMC7 may reduce intracellular exposure of 3-NPA or its reactive intermediates. This combined pattern of metabolic processing and controlled permeability is consistent with adaptive responses to toxic small molecules in Gram-negative bacteria. Importantly, mutations in these same systems in the derived mutants provide a plausible connection between tolerance pathways and the observed increase in kanamycin resistance.

The emergence of aminoglycoside tolerance in the absence of retained Tn5-Mob sequences or canonical acquired resistance determinants further supports a physiological and regulatory interpretation. Perturbations in envelope stress response systems, efflux regulation, and porin expression provide plausible routes for decreased aminoglycoside susceptibility without horizontal acquisition of dedicated resistance genes. These intrinsic resistance mechanisms, arising from regulatory rewiring and altered membrane physiology, are increasingly recognized in environmental and opportunistic bacteria. In the present study, the data suggest that selection under mutagenic and antibiotic conditions can expose linked metabolic and resistance-associated phenotypes, particularly when central metabolism, membrane stress, and transport functions are simultaneously perturbed.

Integration of the LTM01 and LTM14 genomic profiles with the wild-type composite RNA-seq profile showed that several mutant-affected genes belonged to modules transcriptionally engaged during glucose-3NPA growth. These included functions linked to serine metabolism, cysteine pathway entry, one-carbon regulation, nitrogen allocation, amino acid transport, pyruvate processing, cofactor support, and metal-redox homeostasis. This overlap supports the interpretation that impaired 3-NPA utilization in LTM01 and LTM14 is associated with disruption of adaptive support modules rather than loss of a single catabolic entry enzyme. During 3-NPA utilization, EMC7 likely requires coordinated nitrogen redistribution, carbon entry, redox balance, and stress mitigation. Loss-of-function mutations in these connected modules would be expected to reduce the ability of the mutants to assimilate 3-NPA-derived nitrogen, maintain sulfur-redox buffering, process downstream carbon intermediates, and tolerate nitro-redox stress.

Collectively, the results support a candidate model in which 3-NPA utilization in *L. barmai* EMC7 is associated with a distributed metabolic and stress-response framework rather than a strictly linear catabolic pathway (Fig. 3). This framework links carbon entry, nitrogen allocation, cofactor support, sulfur-redox buffering, transport, envelope remodeling, and metal-redox homeostasis. Under this model, the impaired growth of LTM01 and LTM14 may reflect reduced coordination among multiple adaptive processes rather than disruption of a single enzymatic step. This model remains hypothesis-driven, but it provides a useful framework for future functional validation of candidate modules associated with bacterial 3-NPA utilization.

Several caveats should be considered. First, the shared loss-of-function mutations were identified by comparative genome analysis, but individual genetic validation was not performed. Therefore, causality cannot be assigned to any single mutation without targeted complementation or reconstruction of the corresponding variants in the wild-type background. Second, the wild-type composite RNA-seq profile provides pathway-level support for modules engaged during glucose-3NPA growth, but it does not prove the direct biochemical role of each gene in 3-NPA conversion. Future RNA-seq experiments using independently sequenced biological replicates would improve gene-level statistical resolution. Third, enzyme assays, expanded metabolite profiling, or targeted genetic rescue would be required to define the precise biochemical contribution of the candidate nitro-redox, nitrogen allocation, sulfur-redox, and metal-homeostasis modules. Finally, although no stable Tn5- Mob insertion was detected in LTM01 or LTM14 under the sequencing and analysis conditions used, highly fragmented, rearranged, or transient transposon-derived sequences cannot be completely excluded. Therefore, the proposed module-level model should be interpreted as hypothesis-generating rather than as a resolved causal pathway.

Despite these limitations, this study expands the understanding of nitroaliphatic substrate utilization in environmental bacteria by linking 3-NPA depletion, mutant growth impairment, convergent genomic variation, and wild-type composite RNA-seq profiles. The findings identify candidate metabolic and stress-response modules associated with 3-NPA utilization in *L. barmai* EMC7 and provide a basis for future functional validation through complementation, targeted reconstruction, enzyme assays, and metabolite profiling. This perspective is relevant for environmental microbiology, where microbial utilization of toxic or unusual substrates may involve coordinated physiological adaptation rather than single-gene pathway logic.

## 5. CONCLUSION

This study reframes 3-NPA utilization in Leclercia barmai EMC7 as a candidate module-level phenotype rather than a simple single-gene trait. Wild-type EMC7 depleted nearly half of the supplied 3-NPA within 36 h, while the two electroporation-derived mutants, LTM01 and LTM14, showed poor growth under 3-NPA-dependent minimal conditions. Notably, neither mutant retained a detectable stable Tn5-Mob insertion. Instead, both carried convergent loss-of-function mutations in genes linked to nitrogen allocation, carbon-redox metabolism, cofactor-associated processes, transport, and metal-homeostasis functions.

The wild-type composite RNA-seq profile added another layer to this interpretation. Growth under glucose- 3NPA was associated with expression changes in redox enzymes, nitrogen-responsive systems, envelope-stress functions, efflux components, membrane-remodeling genes, and cofactor-linked pathways. Taken together, these observations suggest that 3-NPA utilization in EMC7 may depend on physiological support from several connected modules, not only on a discrete canonical catabolic enzyme. The model remains provisional. Direct tests, including complementation, targeted reconstruction, enzyme assays, and metabolite profiling, will be needed to determine which of these candidate modules have causal roles in 3-NPA utilization.

## Statements and declarations

## Acknowledgements

The authors thank the Department of Biotechnology, University of North Bengal for providing infrastructural support during the work.

## Declaration of generative AI use

During the preparation of this work, the authors used OpenAI ChatGPT (GPT-5.5) to assist with language editing and improvement of overall readability. After using this tool, the authors reviewed and edited the content as needed and take full responsibility for the content of the publication. No experimental data was generated/altered using generative AI.

## CRediT authorship contribution statement

Partha Barman: Data curation, Formal analysis, Investigation, Methodology, Validation, Visualization, Writing – original draft, Writing – review and editing.

Shilpa Sinha: Data curation, Formal analysis, Investigation, Validation, Visualization, Writing – original draft, Writing – review and editing.

Subhajit Sen: Formal analysis, Investigation, Writing – review and editing.

Nibendu Mondal: Data curation, Formal analysis, Investigation, Validation, Writing – review and editing.

Wriddhiman Ghosh: Formal analysis, Resources, Writing – original draft, Writing – review and editing.

Ranadhir Chakraborty: Conceptualization, Methodology, Resources, Supervision, Project administration, Writing – original draft, Writing – review and editing.

All authors read and approved the final manuscript.

## Data Availability Statement

All sequencing data generated in this study are publicly available at NCBI. The EMC7 dataset is available under BioProject <u>PRJNA675190</u>, with SRA <u>SRR15400871</u>, BioSample <u>SAMN16691921</u>, and the Whole Genome Shotgun assembly in GenBank under accession <u>JADMNK000000000</u>. The LTM01 dataset is available under BioProject <u>PRJNA755871</u>, with SRA <u>SRR35965558</u>, BioSample <u>SAMN20845777</u>, and the Whole Genome Shotgun assembly in GenBank under accession <u>JAIZFU000000000</u>. The LTM14 dataset is available under BioProject <u>PRJNA755889</u>, with SRA <u>SRR35965586</u>, BioSample <u>SAMN20845974</u>, and the Whole Genome Shotgun assembly in GenBank under accession <u>JAIKTT000000000</u>. The RNA-seq raw reads generated for wild-type EMC7 under Glu-KNO_3_ control and Glu-3NPA treatment conditions have been deposited in the NCBI Sequence Read Archive under BioProject <u>PRJNA1482400</u>. The Glu-3NPA treatment sample is available under BioSample <u>SAMN61121961</u>, and the Glu-KNO_3_ control sample is available under BioSample <u>SAMN61121962</u>. The corresponding SRA run accessions are <u>SRR39335166</u> and <u>SRR39335167</u> respectively.

## Ethical standards

The present study involved bacterial culture, phenotypic assays, genome analysis, and transcriptome analysis of the previously isolated strain *Leclercia barmai* EMC7. No new animal experimentation, human participants, human-derived samples, or clinical sampling were involved in the present work. The original earthworm-associated isolation of EMC7 was reported previously and was conducted under Institutional Animal Ethics Committee approval, University of North Bengal (Approval no. IAEC/NBU/2022/34). All experiments reported in the present study were conducted in compliance with applicable institutional guidelines and relevant national regulations of India.

---

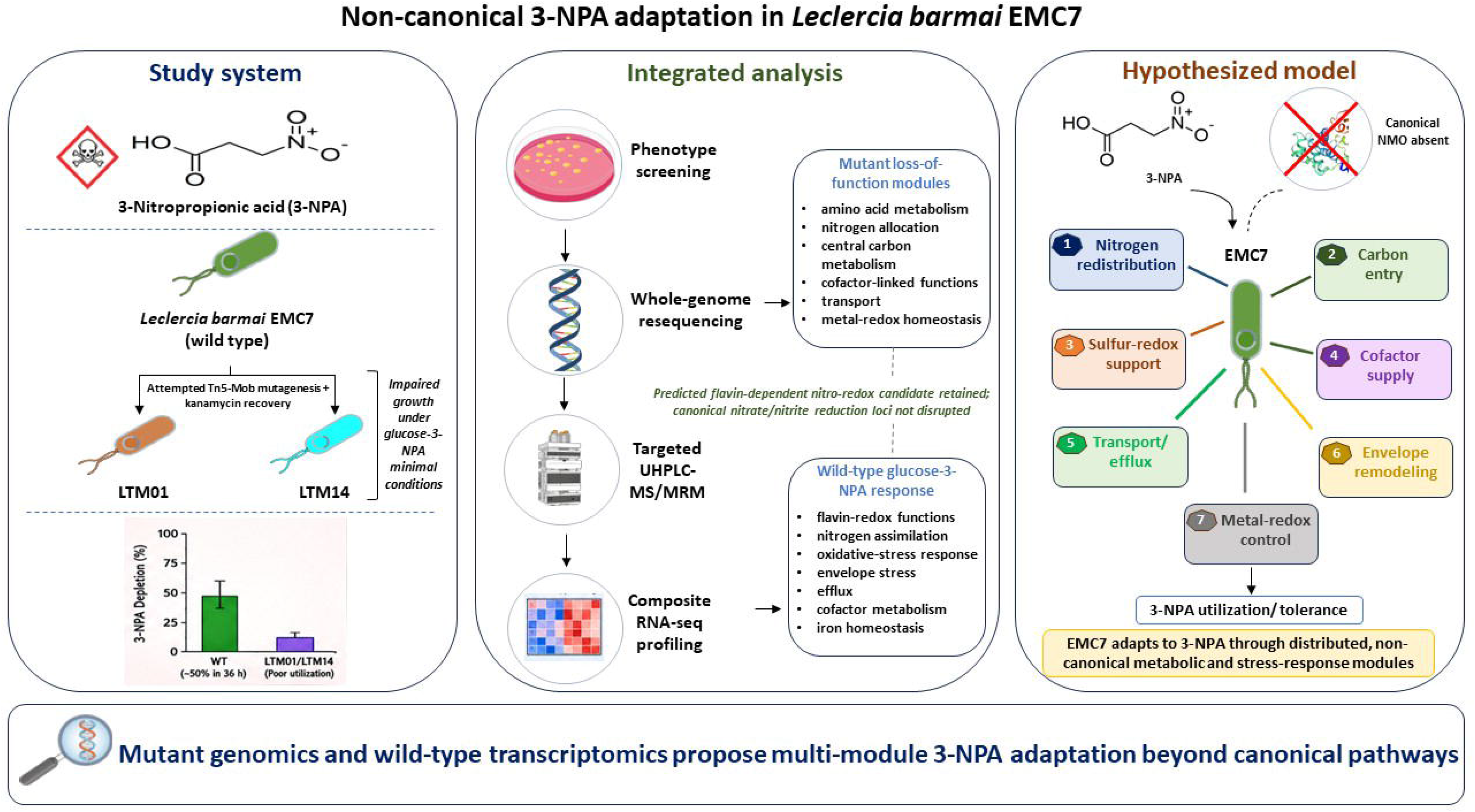

## References

Anderson, R.C., Majak, W., Rassmussen, M.A., Callaway, T.R., Beier, R.C., Nisbet, D.J., Allison, M.J., 2005. Toxicity and Metabolism of the Conjugates of 3-Nitropropanol and 3-Nitropropionic Acid in Forages Poisonous to Livestock. J. Agric. Food Chem. 53, 2344–2350. 10.1021/jf040392j

Bankevich, A., Nurk, S., Antipov, D., Gurevich, A.A., Dvorkin, M., Kulikov, A.S., Lesin, V.M., Nikolenko, S.I., Pham, S., Prjibelski, A.D., Pyshkin, A. V., Sirotkin, A. V., Vyahhi, N., Tesler, G., Alekseyev, M.A., Pevzner, P.A., 2012. SPAdes: A New Genome Assembly Algorithm and Its Applications to Single-Cell Sequencing. Journal of Computational Biology 19, 455–477. 10.1089/cmb.2012.0021

Barman, P., Mondal, N., Sen, S., Chatterjee, S., Sinha, S., Ghosh, W., Chakraborty, R., 2022. Draft Genome Sequences from Two Gram-Negative Bacteria, *Serratia* sp. Strain EWG9 and *Leclercia* sp. Strain EMC7, Isolated from the Earthworm Eisenia fetida. Microbiol. Resour. Announc. 11. 10.1128/mra.00939-21

Barman, P., Sinha, S., Chakraborty, R., 2025. Leclercia barmai sp. nov., isolated from worm castings of Eisenia fetida, is a urease-positive, 3-nitropropionic acid and glycerol-consuming bacterium. Sci. Rep. 15, 5615. 10.1038/s41598-024-78134-7

Basak, C., Mondal, N., Sen, S., Sarkar, J., Ghosh, W., Chakraborty, R., 2021. Draft Genome Sequences of Four Novel Strains of Microbes Isolated from *Lepidocephalichthys guntea*. Microbiol. Resour. Announc. 10. 10.1128/MRA.00621-21

Becker, T., Pasteels, J., Weigel, C., Dahse, H.-M., Voigt, K., Boland, W., 2017. A tale of four kingdoms – isoxazolin-5-one- and 3-nitropropanoic acid-derived natural products. Nat. Prod. Rep. 34, 343–360. 10.1039/C6NP00122J

Bhasin, A., Goryshin, I.Y., Reznikoff, W.S., 1999. Hairpin Formation in Tn5 Transposition. Journal of Biological Chemistry 274, 37021–37029. 10.1074/jbc.274.52.37021

Bilyk, B., Weber, S., Myronovskyi, M., Bilyk, O., Petzke, L., Luzhetskyy, A., 2013. In vivo random mutagenesis of streptomycetes using mariner-based transposon Himar1. Appl. Microbiol. Biotechnol. 97, 351–359. 10.1007/s00253-012-4550-x

Cartman, S.T., Minton, N.P., 2010. A *m ariner* -Based Transposon System for *In Vivo* Random Mutagenesis of *Clostridium difficile*. Appl. Environ. Microbiol. 76, 1103–1109. 10.1128/AEM.02525-09

Connor, C.H., Higgs, C.K., Horan, K., Kwong, J.C., Grayson, M.L., Howden, B.P., Seemann, T., Gorrie, C.L., Sherry, N.L., 2025. Rapid, reference-free identification of bacterial pathogen transmission using optimized split k-mer analysis. Microb. Genom. 11. 10.1099/mgen.0.001347

Diesh, C., Stevens, G.J., Xie, P., De Jesus Martinez, T., Hershberg, E.A., Leung, A., Guo, E., Dider, S., Zhang, J., Bridge, C., Hogue, G., Duncan, A., Morgan, M., Flores, T., Bimber, B.N., Haw, R., Cain, S., Buels, R.M., Stein, L.D., Holmes, I.H., 2023. JBrowse 2: a modular genome browser with views of synteny and structural variation. Genome Biol. 24, 74. 10.1186/s13059-023-02914-z

Field, A., 2024. Discovering statistics using IBM SPSS statistics. Sage publications limited.

Foster, P.L., 2007. Stress-Induced Mutagenesis in Bacteria. Crit. Rev. Biochem. Mol. Biol. 42, 373–397. 10.1080/10409230701648494

Foster, P.L., 2005. Stress responses and genetic variation in bacteria. Mutation Research - Fundamental and Molecular Mechanisms of Mutagenesis 569, 3–11. 10.1016/j.mrfmmm.2004.07.017

Francis, K., Nishino, S.F., Spain, J.C., Gadda, G., 2012. A novel activity for fungal nitronate monooxygenase: Detoxification of the metabolic inhibitor propionate-3-nitronate. Arch. Biochem. Biophys. 521, 84–89. 10.1016/j.abb.2012.03.015

Francis, K., Smitherman, C., Nishino, S.F., Spain, J.C., Gadda, G., 2013. The biochemistry of the metabolic poison propionate 3 nitronate and its conjugate acid, 3 nitropropionate. IUBMB Life 65, 759–768. 10.1002/iub.1195

Frenoy, A., Bonhoeffer, S., 2018. Death and population dynamics affect mutation rate estimates and evolvability under stress in bacteria. PLoS Biol. 16, e2005056. 10.1371/journal.pbio.2005056

Galhardo, R.S., Hastings, P.J., Rosenberg, S.M., 2007. Mutation as a Stress Response and the Regulation of Evolvability. Crit. Rev. Biochem. Mol. Biol. 42, 399–435. 10.1080/10409230701648502

Geddes, J.W., Bondada, V., Pang, Z., 2000. Mechanisms of 3-Nitropropionic Acid Neurotoxicity, in: Mitochondrial Inhibitors and Neurodegenerative Disorders. Humana Press, Totowa, NJ, pp. 107–120. 10.1007/978-1-59259-692-8_7

Hamilton, B.F., Gould, D.H., Gustine, D.L., 2000. History of 3-Nitropropionic Acid, in: Mitochondrial Inhibitors and Neurodegenerative Disorders. Humana Press, Totowa, NJ, pp. 21–33. 10.1007/978-1-59259-692-8_2

Huangyang, P., Simon, M.C., 2018. Hidden features: exploring the non-canonical functions of metabolic enzymes. Dis. Model. Mech. 11. 10.1242/dmm.033365

Jilk, R.A., Makris, J.C., Borchardt, L., Reznikoff, W.S., 1993. Implications of Tn5-associated adjacent deletions. J. Bacteriol. 175, 1264–1271. 10.1128/jb.175.5.1264-1271.1993

Johnson, M., Zaretskaya, I., Raytselis, Y., Merezhuk, Y., McGinnis, S., Madden, T.L., 2008. NCBI BLAST: a better web interface. Nucleic Acids Res. 36, W5–W9. 10.1093/nar/gkn201

Kivisaar, M., 2003. Stationary phase mutagenesis: mechanisms that accelerate adaptation of microbial populations under environmental stress. Environ. Microbiol. 5, 814–827. 10.1046/j.1462-2920.2003.00488.x

Li, N., Jin, K., Bai, Y., Fu, H., Liu, L., Liu, B., 2020. Tn5 Transposase Applied in Genomics Research. Int. J. Mol. Sci. 21, 8329. 10.3390/ijms21218329

Matic, I., 2017. Molecular mechanisms involved in the regulation of mutation rates in bacteria. Period. Biol. 118. 10.18054/pb.v118i4.4601

Mukhopadhyaya, P.N., Deb, C., Lahiri, C., Roy, P., 2000. A *soxA* Gene, Encoding a Diheme Cytochrome *c*, and a *sox* Locus, Essential for Sulfur Oxidation in a New Sulfur Lithotrophic Bacterium. J. Bacteriol. 182, 4278–4287. 10.1128/JB.182.15.4278-4287.2000

Nishino, S.F., Shin, K.A., Payne, R.B., Spain, J.C., 2010. Growth of Bacteria on 3-Nitropropionic Acid as a Sole Source of Carbon, Nitrogen, and Energy. Appl. Environ. Microbiol. 76, 3590–3598. 10.1128/AEM.00267-10

Reese, M.G., 2001. Application of a time-delay neural network to promoter annotation in the Drosophila melanogaster genome. Comput. Chem. 26, 51–56. 10.1016/S0097-8485(01)00099-7

Reynolds, N.C., Lin, W., 2000. The Neurochemistry of 3-Nitropropionic Acid, in: Mitochondrial Inhibitors and Neurodegenerative Disorders. Humana Press, Totowa, NJ, pp. 35–49. 10.1007/978-1-59259-692-8_3

Reznikoff, W.S., 2003. Tn *5* as a model for understanding DNA transposition. Mol. Microbiol. 47, 1199–1206. 10.1046/j.1365-2958.2003.03382.x

Rogowska-van der Molen, M.A., Nagornîi, D., Coolen, S., de Graaf, R.M., Berben, T., van Alen, T., Janssen, M.A.C.H., Rutjes, F.P.J.T., Jansen, R.S., Welte, C.U., 2022. Insect Gut Isolate Pseudomonas sp. Strain Nvir Degrades the Toxic Plant Metabolite Nitropropionic Acid. Appl. Environ. Microbiol. 88. 10.1128/aem.00719-22

Saha, T., Ranjan, V.K., Ganguli, S., Thakur, S., Chakraborty, B., Barman, P., Ghosh, W., Chakraborty, R., 2019. Pradoshia eiseniae gen. nov., sp. nov., a spore-forming member of the family Bacillaceae capable of assimilating 3-nitropropionic acid, isolated from the anterior gut of the earthworm Eisenia fetida. Int. J. Syst. Evol. Microbiol. 69, 1265–1273. 10.1099/ijsem.0.003304

Scallet, A.C., Haley, R.L., Scallet, D.M., Duhart, H.M., Binienda, Z.K., 2003. 3 Nitropropionic Acid Inhibition of Succinate Dehydrogenase (Complex II) Activity in Cultured Chinese Hamster Ovary Cells. Ann. N. Y. Acad. Sci. 993, 305–312. 10.1111/j.1749-6632.2003.tb07538.x

Sen, S., Mondal, N., Ghosh, W., Chakraborty, R., 2022. Inducible boron resistance via active efflux in Lysinibacillus and Enterococcus isolates from boron-contaminated agricultural soil. BioMetals 35, 215–228. 10.1007/s10534-021-00359-0

Sen, S., Saha, T., Bhattacharya, S., Nidhi, Mondal, N., Ghosh, W., Chakraborty, R., 2020. Draft Genome Sequences of Two Boron-Tolerant, Arsenic-Resistant, Gram-Positive Bacterial Strains, Lysinibacillus sp. OL1 and Enterococcus sp. OL5, Isolated from Boron-Fortified Cauliflower-Growing Field Soils of Northern West Bengal, India. Microbiol. Resour. Announc. 9. 10.1128/MRA.01438-19

Simon, R., Priefer, U., Pühler, A., 1983. A Broad Host Range Mobilization System for In Vivo Genetic Engineering: Transposon Mutagenesis in Gram Negative Bacteria. Bio/Technology 1, 784–791. 10.1038/nbt1183-784

Snaebjornsson, M.T., Schulze, A., 2018. Non-canonical functions of enzymes facilitate cross-talk between cell metabolic and regulatory pathways. Exp. Mol. Med. 50, 1–16. 10.1038/s12276-018-0065-6

Upadhayay, S., Yedke, N.G., Rahi, V., Singh, S., Kumar, S., Arora, A., Chandolia, P., Kaur, P., Kumar, M., Koshal, P., Jamwal, S., Kumar, P., 2023. An Overview of the Pathophysiological Mechanisms of 3- Nitropropionic Acid (3-NPA) as a Neurotoxin in a Huntington’s Disease Model and Its Relevance to Drug Discovery and Development. Neurochem. Res. 48, 1631–1647. 10.1007/s11064-023-03868-1

Wall, J.D., Murnan, T., Argyle, J., English, R.S., Rapp-Giles, B.J., 1996. Transposon mutagenesis in Desulfovibrio desulfuricans: development of a random mutagenesis tool from Tn7. Appl. Environ. Microbiol. 62, 3762–3767. 10.1128/aem.62.10.3762-3767.1996

Wright, B.E., 2004. Stress directed adaptive mutations and evolution. Mol. Microbiol. 52, 643–650. 10.1111/j.1365-2958.2004.04012.x

Zhang, Y., Xu, S., Chai, C., Yang, S., Jiang, W., Minton, N.P., Gu, Y., 2016. Development of an inducible transposon system for efficient random mutagenesis in *Clostridium acetobutylicum*. FEMS Microbiol. Lett. 363, fnw065. 10.1093/femsle/fnw065

