## Supplemental Table for "Convergent genomic variation and wild-type composite RNA-seq profiling identify candidate modules associated with 3-nitropropionic acid utilization in *Leclercia barmai*"

### Electronic Supplementary Material

**SI 1** GYT medium

| Composition | Amount per litre |
|---|---|
| Tryptone | 2.5 g |
| Yeast extract | 1.25 g |
| Glycerol | 100 mL |

**SI 2** SOC medium

| Composition | Amount per litre |
|---|---|
| Tryptone | 20 g |
| Yeast extract | 5 g |
| NaCl | 0.5 g |
| KCl | 0.186 g |
| $MgCl_2$ | 10mM |
| $MgSO_4$ | 10mM |
| Glucose | 3.6 g |

**SI 3** SOB medium

| Composition | Amount per litre |
|---|---|
| Tryptone | 20 g |
| Yeast extract | 5 g |
| NaCl | 0.5 g |
| KCl | 0.186 g |
| $MgCl_2$ | 10 mM |
| $MgSO_4$ | 10 mM |
| Agar | 15 g |

**SI 4** Screening of kan$^r$ colonies (generated following electroporation of pSUP5011:Tn5-Mob into EMC7) for mutants impaired in 3-NPA utilization

All kanamycin-resistant colonies were picked up individually with sterile toothpicks and spotted in the middle of each numbered (serially from 1 to 1009) grid of a total of 25 LA-kanamycin (0.04 g/L) plates (Plate A–Plate Y). From this pool, 40 colonies, randomly selected using a random number generator (randomization being carried out without replacement; https://www.randomizer.org), were picked to construct an LA-kanamycin master plate. The master plate was incubated overnight for colony growth (of 40 colonies picked, 39 manifested growth) before being replica-plated onto fresh agar plates composed of mineral salts medium (MSM) supplemented with glucose, $KNO_3$, and kanamycin or MSM supplemented with glucose, 3- NPA and kanamycin or MSM supplemented with 3-NPA or Luria broth supplemented with kanamycin or Luria broth (SI 5 and SI 6; kan used: 0.04 g/L). Following 24 h incubation of the replica plates at 30 °C, two colonies that did not show growth on plates composed of MSM supplemented with glucose, 3-NPA and kanamycin or MSM supplemented with 3-NPA but grew on plates composed of MSM supplemented with glucose, $KNO_3$, and kanamycin or LA supplemented with kanamycin or LA were clonally purified on MSM supplemented with glucose, $KNO_3$, and kanamycin plates (SI 5; kanamycin used: 0.04 g/L) for further analysis.

**SI 5** Minimal salt medium supplemented with Glucose as carbon source and $KNO_3$ as sole nitrogen source

| **Composition** | **Amount per litre** |
|---|---|
| $KH_2PO_4$ | 3 g |
| $Na_2HP0_4$ | 6 g |
| NaCl | 5 g |
| $MgSO_4$ | 0.1 g |
| Glucose | 8 g |
| $KNO_3$ | 0.875 g |

**SI 6** Minimal salt medium supplemented with Glucose as carbon source and 3-NPA as sole nitrogen source

| **Composition** | **Amount per litre** |
|---|---|
| $KH_2PO_4$ | 3 g |
| $Na_2HP0_4$ | 6 g |
| NaCl | 5 g |

| | |
|---|---|
| $MgSO_4$ | 0.1 g |
| Glucose | 8 g |
| 3-NPA | 0.06 g |

**SI 7** Minimal salt medium supplemented with 3-NPA as sole carbon and nitrogen source

| Composition | Amount per litre |
|---|---|
| $KH_2PO_4$ | 3 g |
| $Na_2HP0_4$ | 6 g |
| NaCl | 5 g |
| $MgSO_4$ | 0.1 g |
| 3-NPA | 0.12 g |

**SI 8** Minimal salt medium supplemented with Glucose as carbon source, $KNO_3$ as sole nitrogen source, kanamycin and PAβN

| Composition | Amount per litre |
|---|---|
| $KH_2PO_4$ | 3 g |
| $Na_2HP0_4$ | 6 g |
| NaCl | 5 g |
| $MgSO_4$ | 0.1 g |
| Glucose | 8 g |
| $KNO_3$ | 0.875 g |
| Kanamycin | 0.04 g |
| PaβN (when used) | 0.1 g |

**SI 9** Functional overview of the DEG response

| **Functional class** | **DEGs** | **Up** | **Down** | **Interpretation** |
|---|---|---|---|---|
| Direct nitro/flavin oxygenation or reduction | **40** | **27** | 13 | Strongest evidence for noncanonical NPA transformation chemistry |
| Nitrate/nitrite sensing, nitrate reduction, Mo cofactors | **10** | **7** | 3 | NPA condition activates inorganic nitrogen handling despite KNO3 control |
| Nitrogen assimilation/regulation | **21** | **15** | 6 | NPA-derived nitrogen is being processed, but full ammonium assimilation is not proven |
| Carbon entry/central metabolism | **111** | **62** | 49 | Carbon flux is remodelled, but glucose background weakens carbon-use claims |
| Redox, respiration, quinone modules | **93** | **60** | 33 | Electron transport is rewired under NPA stress |

| | | | | |
|---|---|---|---|---|
| Oxidative stress, Fe-S, DNA repair | 21 | 8 | 13 | Selective peroxide and Fe-S repair response, not a broad antioxidant response |
| Envelope, permeability, efflux | 75 | 30 | 45 | Strong tolerance phenotype, reduced influx and increased export |
| Regulators, chaperones, stress proteins | 126 | 48 | 78 | Global transcriptional reprogramming |

**SI 10** Upregulated profile relevant to 3-NPA metabolism

| Process | Upregulated genes | logFC | Interpretation |
|---|---|---|---|
| Nitro compound reduction | oxygen-insensitive NAD(P)H nitroreductase, ITX56_11945 | +0.654 | Direct candidate for nitro-group redox handling. Bacterial oxygen-insensitive nitroreductases are FMN-linked NAD(P)H enzymes that reduce nitro groups through two-electron chemistry. |
| Flavin-dependent oxidation | flavin-dependent monooxygenase, ITX56_03450 | +0.942 | Candidate for oxidative 3-NPA/P3N activation. Not proof of NMO activity. |
| FMN monooxygenase-like activity | putative FMN-dependent luciferase-like monooxygenase, ITX56_04510 | +0.768 | Candidate oxygenating enzyme. Supports noncanonical flavin route. |
| Luciferase-like monooxygenase | luciferase-like monooxygenase, ITX56_10100 | +0.613 | Secondary monooxygenase candidate. |
| LLM flavin oxidoreductases | ITX56_17880, ITX56_18085 | +1.030, +0.813 | Important because Nvir used a flavin oxidoreductase-like noncanonical enzyme for NPA degradation. |
| Flavin recycling | flavin reductase family proteins, ITX56_14305, ITX56_17785 | +0.922, +0.915 | Supplies reduced flavin to monooxygenase-like reactions. |
| FMN reduction | NADPH-dependent FMN reductase, ITX56_16560 | +0.663 | Supports reduced FMN pool. |

| | | | |
|---|---|---|---|
| NADH-flavin oxidation | NADH:flavin oxidoreductase/NADH oxidase, ITX56_07650 | +0.603 | Links NADH oxidation with flavin redox cycling. |
| FAD oxidoreduction | FAD-binding oxidoreductases, ITX56_18255, ITX56_10560 | +0.898, +0.690 | Broad redox support. |
| Nitrate/nitrite sensing | NarX, ITX56_03780 | +0.697 | NarX is a nitrate/nitrite-sensing histidine kinase. Its induction supports sensing of NPA-derived nitrite/nitrate or altered nitrogen redox state. (MDPI) |
| Nitrate reduction | nitrate reductase, ITX56_03765, nitrate reductase alpha, ITX56_04405 | +0.675, +0.605 | Supports nitrate processing under NPA condition. |
| Nitrate reductase maturation | nitrate reductase Mo-cofactor assembly chaperone, ITX56_04395 | +0.818 | Supports assembly of molybdo-nitrate reductase system. |
| Molybdate acquisition | molybdate ABC transporter substrate-binding protein, ITX56_10985 | +1.208 | Supports Mo-dependent nitrate reductase activity. |
| MoCo biosynthesis | molybdenum cofactor biosynthesis protein B, ITX56_13605 | +0.886 | Reinforces nitrate reductase readiness. |
| Nitrogen regulation | nitrogen regulation protein NR(I), ITX56_21235 | +0.590 | Indicates nitrogen-status response. |
| Glutamine synthetase regulation | GlnE-like adenylyltransferase, ITX56_10705 | +0.587 | Links nitrogen assimilation with GS control. |
| Carbon skeleton adjustment | 3-oxoadipate enol-lactonase, ITX56_03505 | +1.256 | Suggests activation of downstream organic-acid/aromatic-acid metabolism. Not direct proof of 3OP use. |
| Lactate/redox carbon branch | FMN-dependent L-lactate dehydrogenase LldD, ITX56_19865 | +0.666 | Suggests redox balancing through organic acid metabolism. |

| | | | |
|---|---|---|---|
| NADH respiration | NADH-quinone oxidoreductase NuoB, ITX56_12190 | +1.563 | Strong signal for respiratory NADH oxidation. |
| Na+-NADH:ubiquinone reductase | ITX56_20685 | +0.720 | Adds alternative NADH-to-quinone electron flow. |
| Peroxide defense | manganese catalase family protein, ITX56_12930 | +0.846 | Consistent with H2O2 burden expected from oxidative NPA/P3N chemistry. |
| Thiol detox | glutathione transferase, ITX56_01145 | +0.782 | Supports electrophile/redox detox. |
| Glutathione/cysteine export or redox transport | CydC, ITX56_16300 | +0.764 | Supports envelope and redox protection. |
| Thioredoxin family protein | ITX56_18315 | +0.783 | Supports protein thiol repair. |
| Fe-S repair | SufD, ITX56_06155 | +0.867 | SUF is commonly induced under oxidative stress or iron limitation in bacteria. |
| DNA repair | DNA repair protein, ITX56_10430 | +0.709 | Supports damage-response activation. |
| Envelope stress | CpxP, ITX56_21000 | +2.131 | Major envelope-stress signal. CpxP participates in Cpx envelope stress control and misfolded-protein handling. |
| Acid/toxic stress | YqgB, ITX56_19185 | +2.605 | Strong stress marker. |
| Membrane stress | YncL, ITX56_06355 and ITX56_06855 | +1.403, +0.948 | Supports membrane stress adaptation. |
| Efflux | RND permease, ITX56_18475, RND adaptor, ITX56_18470, RND permease, ITX56_11995, MdtC, ITX56_02000 | +1.063, +0.785, +0.746, +0.629 | Supports active extrusion of toxic compound or toxic byproducts. Gram-negative envelope permeability and efflux jointly reduce intracellular compound accumulation. |
| TolC-type outer membrane channel | ITX56_13045 | +0.864 | Fits RND-TolC efflux architecture. |

**SI 11** Downregulated profile relevant to tolerance and metabolic rerouting

| Process | Downregulated genes | logFC | Interpretation |
|---|---|---|---|
| Major porin shutdown | OmpC, ITX56_01435 | -4.521 | Strong reduction in outer-membrane influx route. |
| Outer-membrane structure | OmpA, ITX56_16625 | -3.729 | Reduced permeability and envelope restructuring. Porins regulate passive transport and envelope integrity. |
| Sugar/nutrient porin | maltoporin, ITX56_09385 | -3.337 | Reduced carbohydrate uptake routes, consistent with stress-state narrowing of permeability. |
| Other porins | ITX56_14415, ITX56_21145, ITX56_18895, ITX56_16495 | -1.970, -1.805, -1.518, -0.816 | Broad porin repression. |
| Lipoprotein/envelope proteins | Lpp, ITX56_06135, OsmE, ITX56_06580 | -3.451, -3.672 | Envelope remodeling under toxic condition. |
| LPS/capsule-related genes | LPS biosynthesis protein, ITX56_02210, capsular polysaccharide protein, ITX56_07830 | -3.577, -4.556 | Surface architecture is strongly remodeled. |
| Stationary phase/ribosome hibernation | RaiA, ITX56_13365, RMF, ITX56_16605, HPF-like, ITX56_09875 | -6.939, -3.232, -2.267 | EMC7 is not entering a simple dormancy program. It stays metabolically responsive. |
| Universal stress proteins | UspG, ITX56_14320, UspA, ITX56_22090, UspC, ITX56_02835 | -3.000, -2.688, -1.630 | Stress response is selective, not a broad USP-dominated response. |
| Succinate oxidation | SdhB, ITX56_13900, Sdh cytochrome b556 subunit, ITX56_13915 | -0.718, -1.016 | Suggests TCA/respiratory branch suppression. This is relevant because 3-NPA is known for succinate dehydrogenase toxicity in eukaryotes. |

| | | | |
|---|---|---|---|
| Cytochrome bd branch | CydX, ITX56_13850, YbgE, ITX56_13845 | -1.984, -1.148 | Respiratory branch shift, not uniform respiration induction. |
| Antioxidant genes down | Mn-SOD, ITX56_21020, Cu-Zn SOD, ITX56_05890, TrxA, ITX56_22690, thiol peroxidase, ITX56_00520 | -1.049, -0.593, -0.935, -0.751 | The oxidative response is not general Sox/SOD-like. Catalase, GST, SufD, and thioredoxin-family genes are more relevant here. |
| Fe-S genes down | IscX, ITX56_00295, NfuA, ITX56_22515, SufE, ITX56_06145 | -1.359, -0.660, -0.811 | Fe-S maintenance is reorganized, with SufD up but other Fe-S components down. |
| Efflux gene down | HlyD family efflux adaptor, ITX56_19680 | -1.579 | Efflux response is selective, not all pumps increase. |
| Sugar metabolism down | glycerol-3-phosphate dehydrogenase, ITX56_22555, GAPDH, ITX56_06730, fructose PTS IIB, ITX56_20445 | -2.755, -1.133, -2.769 | Carbon flux is redirected away from some routine carbohydrate modules. |

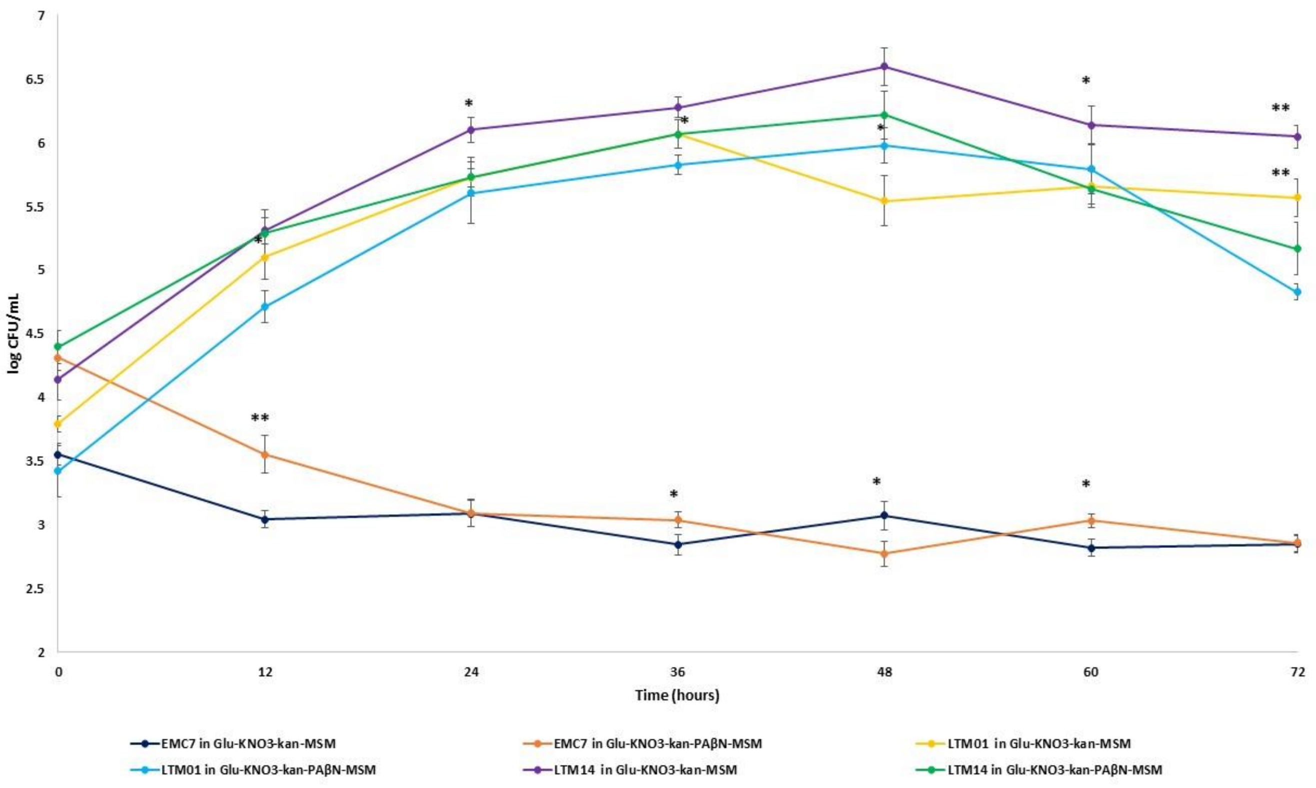

**Supplementary Figure S1. Effect of PAβN on growth of EMC7, LTM01, and LTM14 in MSM supplemented with Glucose, KNO3, and kanamycin in presence and absence of PAβN: The Y-axis represent mean log CFU/mL.** The X-axis represents time of incubation in hours. The error bars represent SD from three replicates. A three-way ANOVA showed significant effects of strain, PAβN condition, time, and strain × condition × time interaction on log CFU/mL. The asterisks indicate within-strain comparison between Glucose-$KNO_3$-kanamycin-MSM and Glucose-$KNO_3$-kanamycin-MSM + PAβN at the same time point. ns, $p \geq 0.05$; *, $p < 0.05$; **, $p < 0.01$; ***, $p < 0.001$.

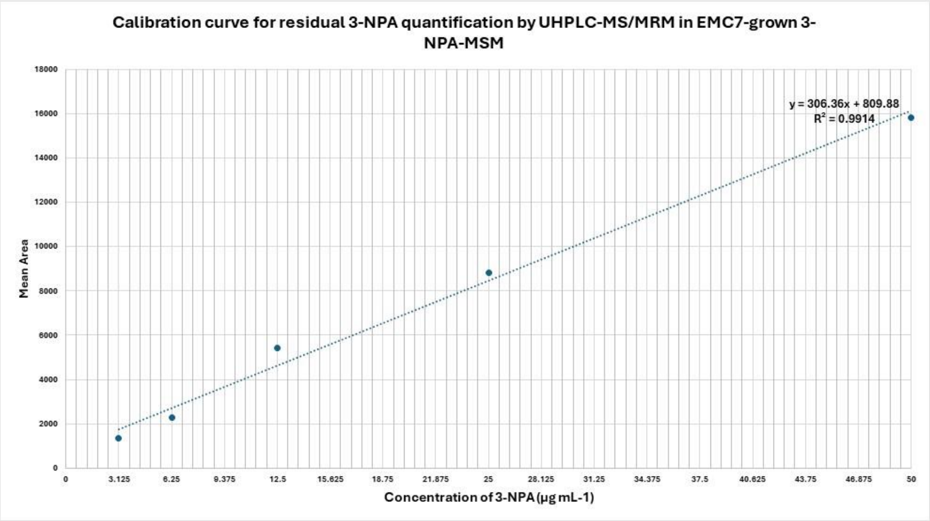


**Supplementary Figure S2a:** Calibration curve for UHPLC-MS/MRM-based quantification of 3-NPA. Mean peak area was plotted against 3-NPA standard concentration, 3.125 to 50 mg $L^{-1}$. Linear regression showed $y = 306.36x + 809.88$ with $R^2 = 0.9914$.

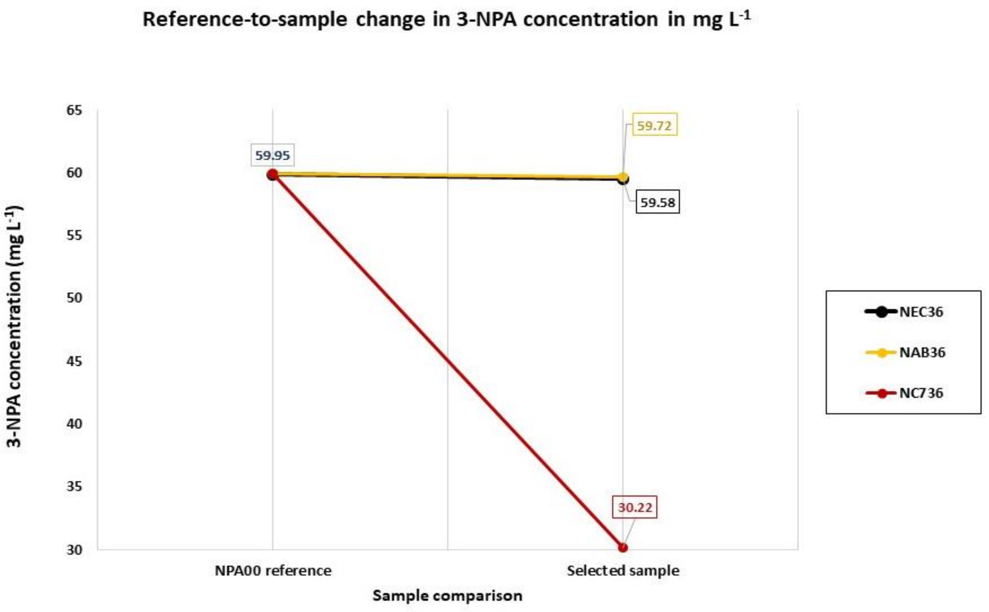


**Supplementary figure S2b.** Reference-to-sample change in residual 3-NPA concentration. Residual 3-NPA in NPA00, the 0 h reference supernatant, was 59.95 mg $L^{-1}$. After 36 h, *E.coli* K12 at 36 h (NEC36) and abiotic control at 36 h (NAB36) retained near-reference concentrations, 59.58 and 59.72 mg $L^{-1}$, respectively, indicating minimal non-biological loss. In contrast, EMC7 at 36 h (NC736) decreased to 30.22 mg $L^{-1}$, corresponding to 49.6% depletion of parent 3-NPA during EMC7 incubation.

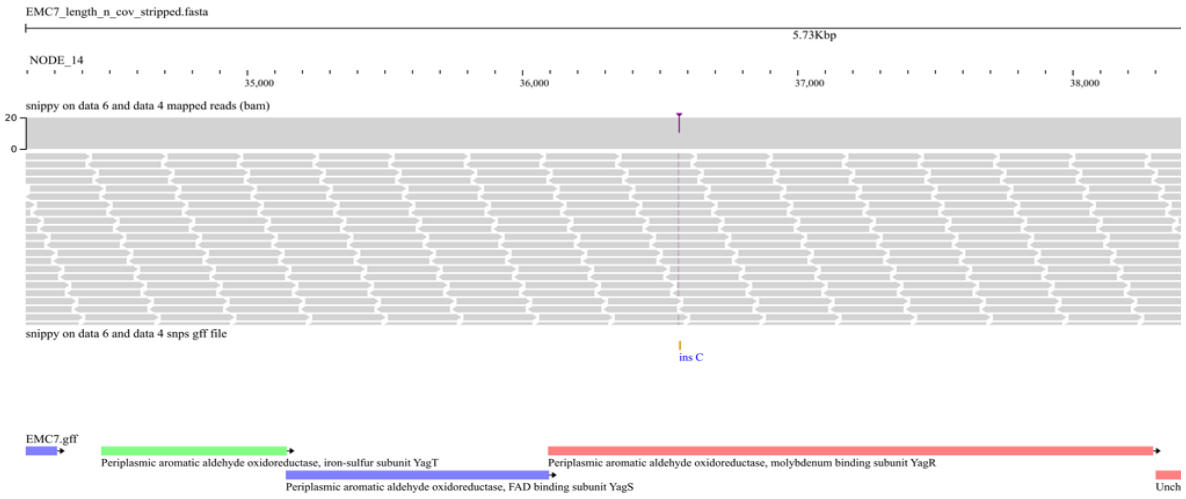

**Supplementary Figure S3a:** ins C mutation in Periplasmic aromatic aldehyde oxidoreductase, molybdenum binding subunit *yagR* gene in LTM01.

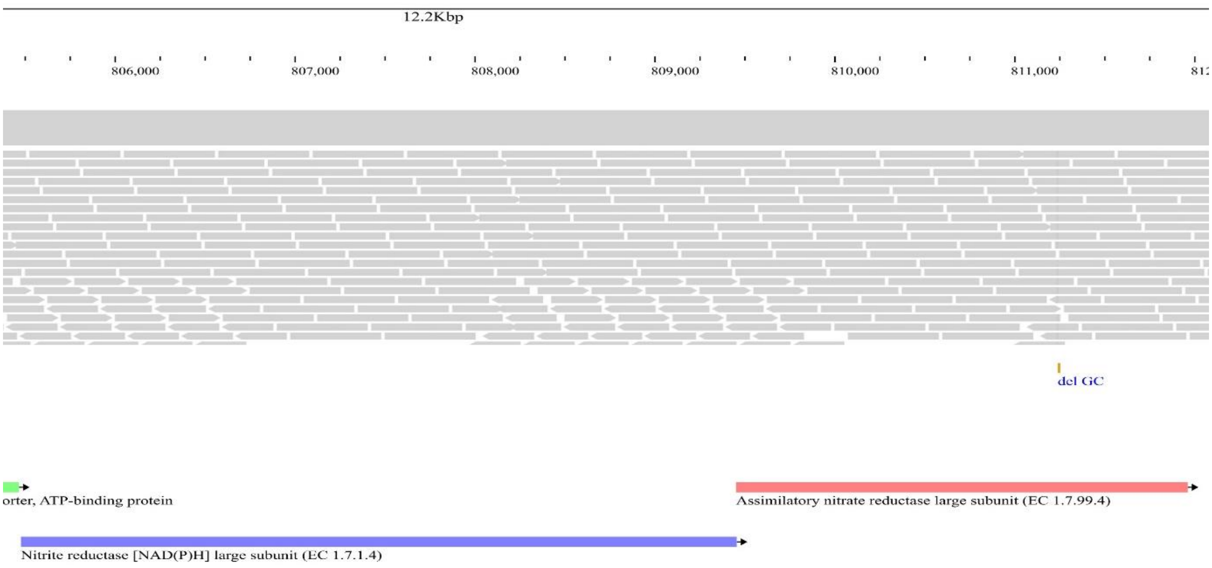


**Supplementary Figure S3b:** del GC mutation in Assimilatory nitrate reductase (large subunit) gene *nasA* in LTM01.

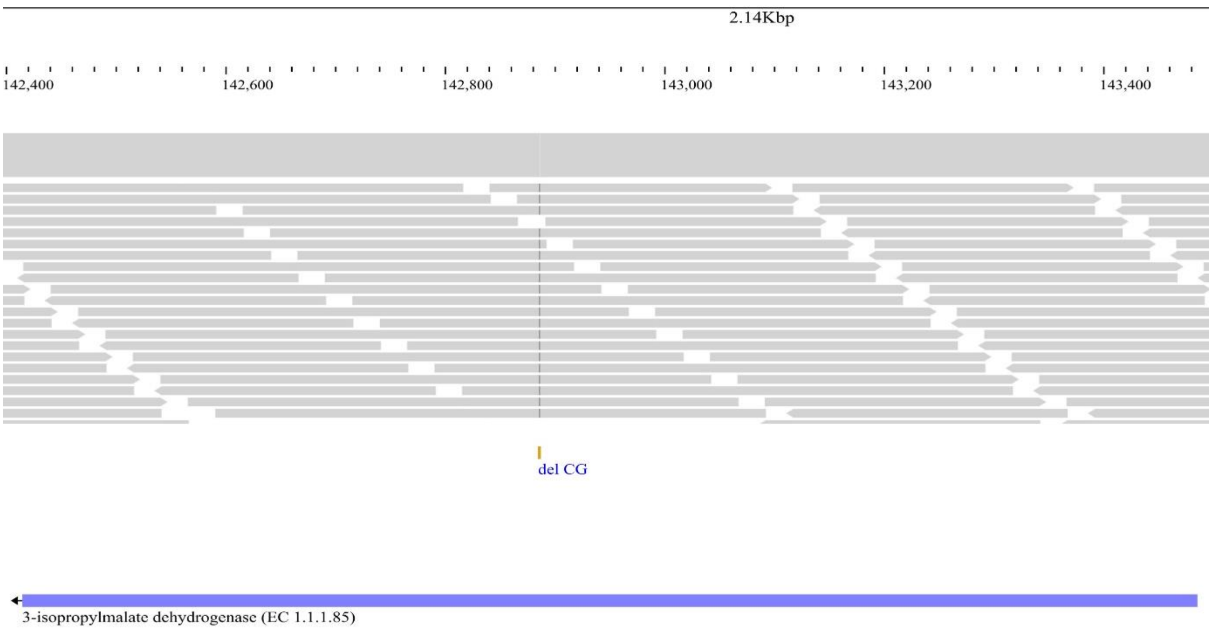


**Supplementary Figure S3c:** del CG mutation in 3-isopropylmalate dehydrogenase gene *leuB* in LTM01.

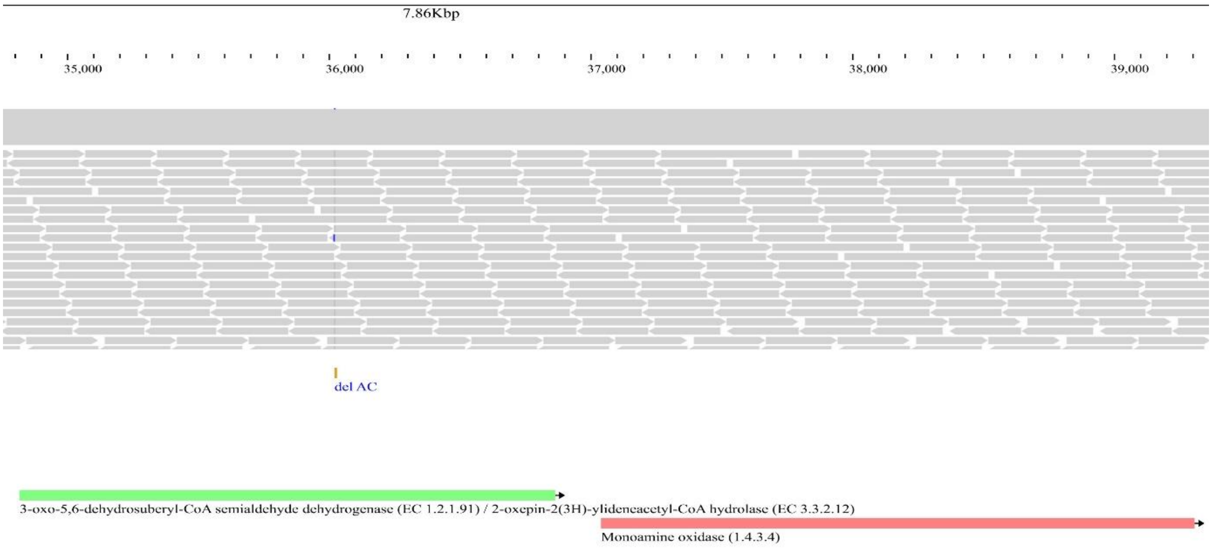


**Supplementary Figure S3d:** del AC mutation in 3-oxo-5,6-dehydrosuberyl-CoA semialdehyde dehydrogenase gene *paaZ* in LTM01.

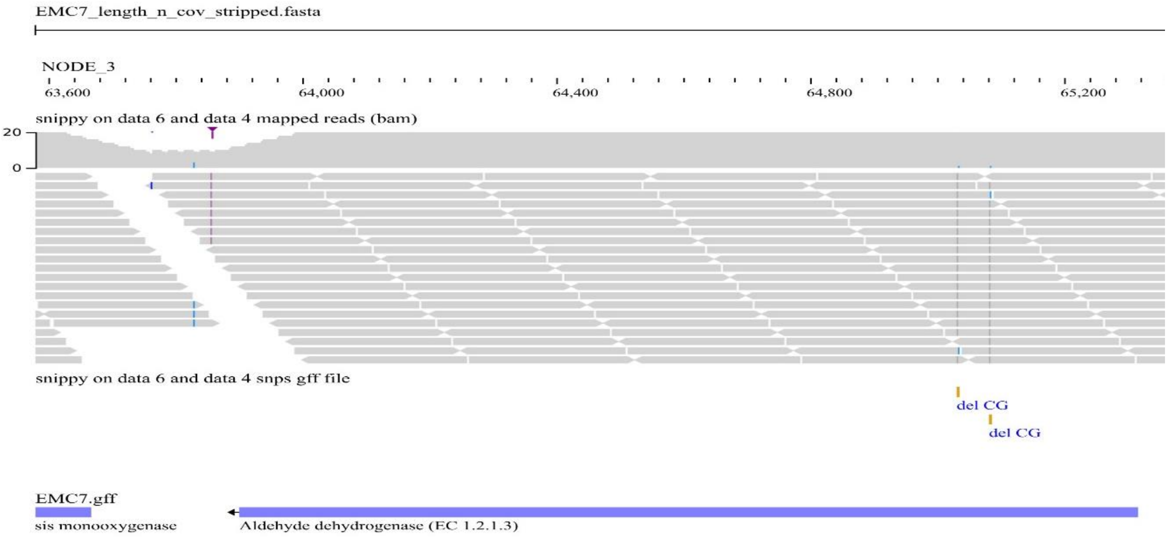


**Supplementary Figure S3e:** Multiple deletion mutations (del CG, del CG) in Aldehyde dehydrogenase gene *aldB* in LTM01.

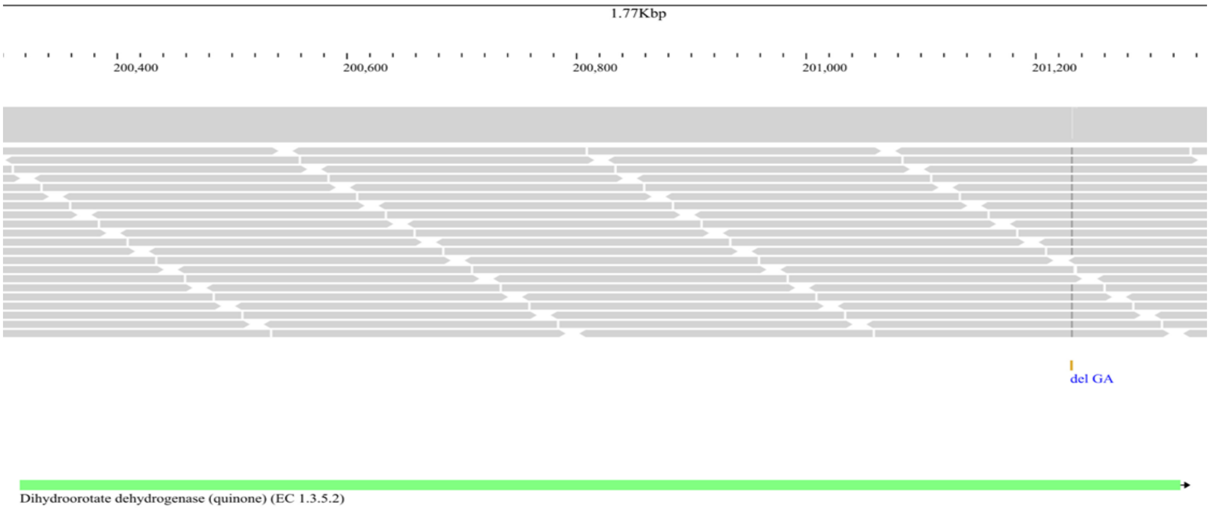


**Supplementary Figure S3f:** del GA mutation in Dihydroorotate dehydrogenase (quinone) gene *pyrD* in LTM01.

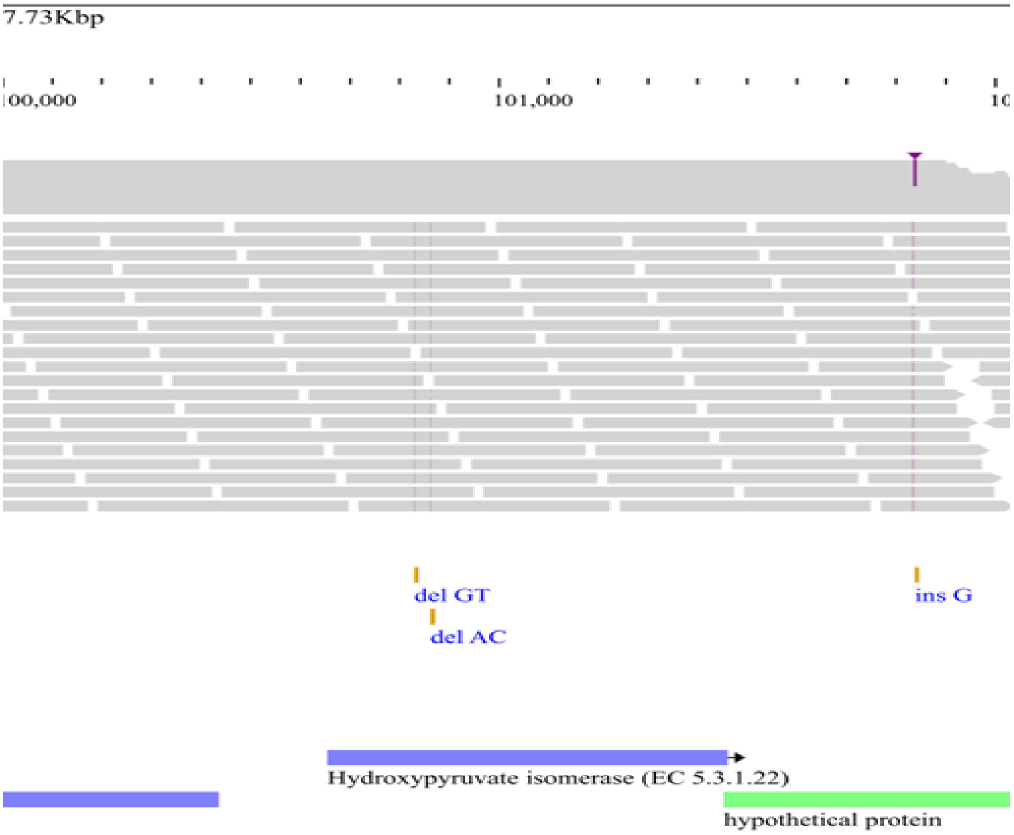


**Supplementary Figure S3g:** Multiple deletion mutations (del GT, del AC) in Hydroxypyruvate isomerase gene *hyi* in LTM01.

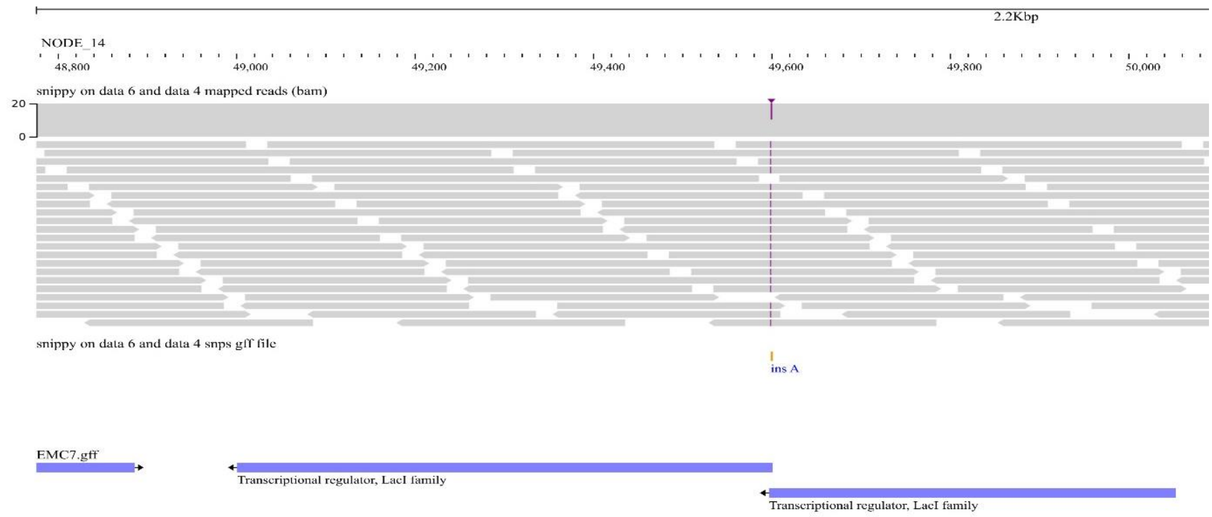


**Supplementary Figure S4a:** ins A mutation in Transcriptional regulator gene, *lacI* family in LTM01.

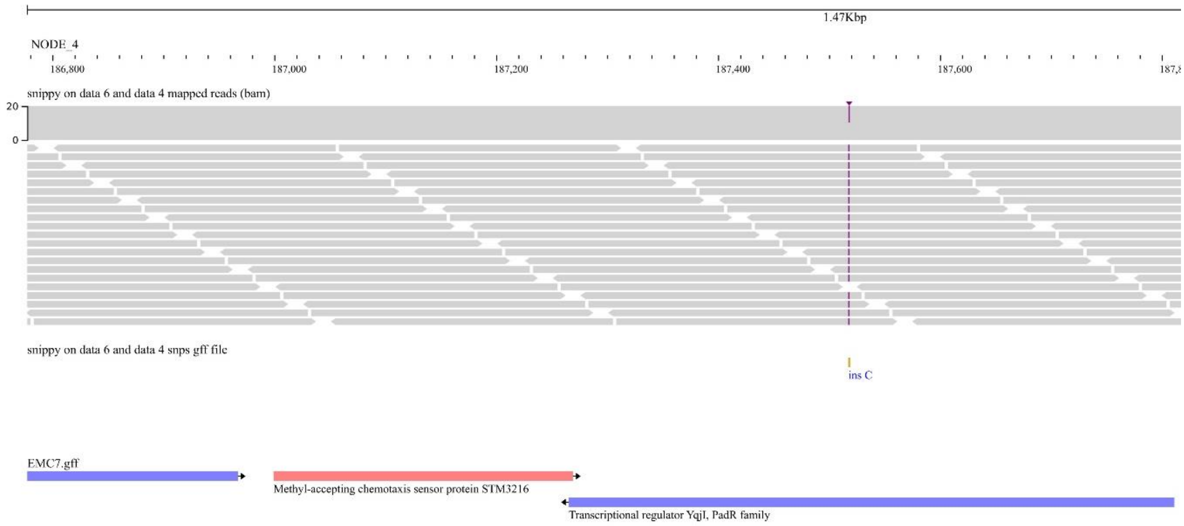


**Supplementary Figure S4b:** ins C mutation in Transcriptional regulator gene *yqjI*, *padR* family in LTM01.

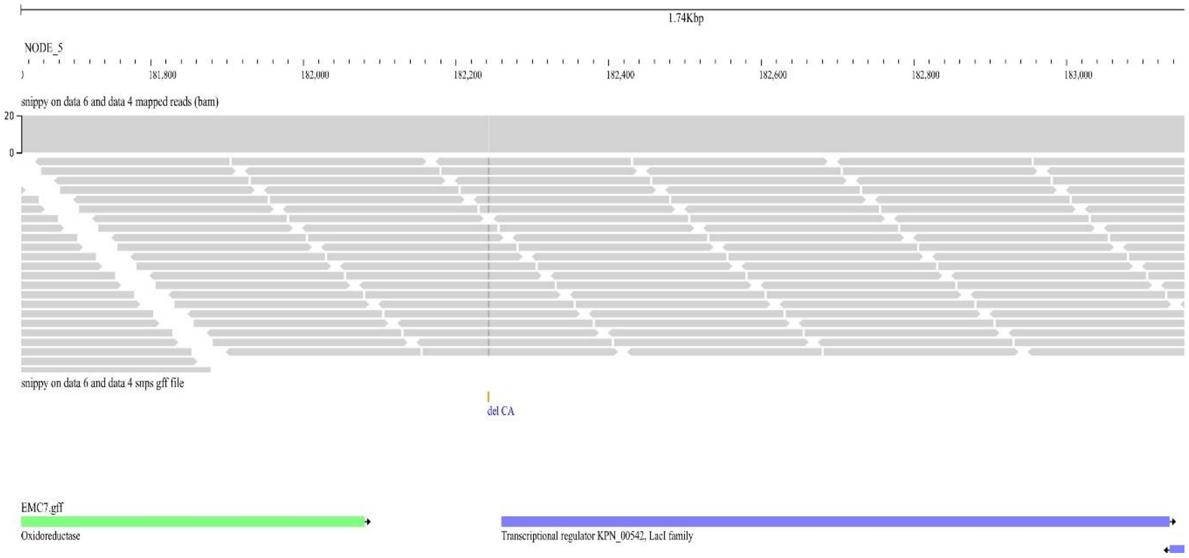


**Supplementary Figure S5:** del CA mutation in Transcriptional regulator KPN_00542 gene, *lacI* family in LTM14.

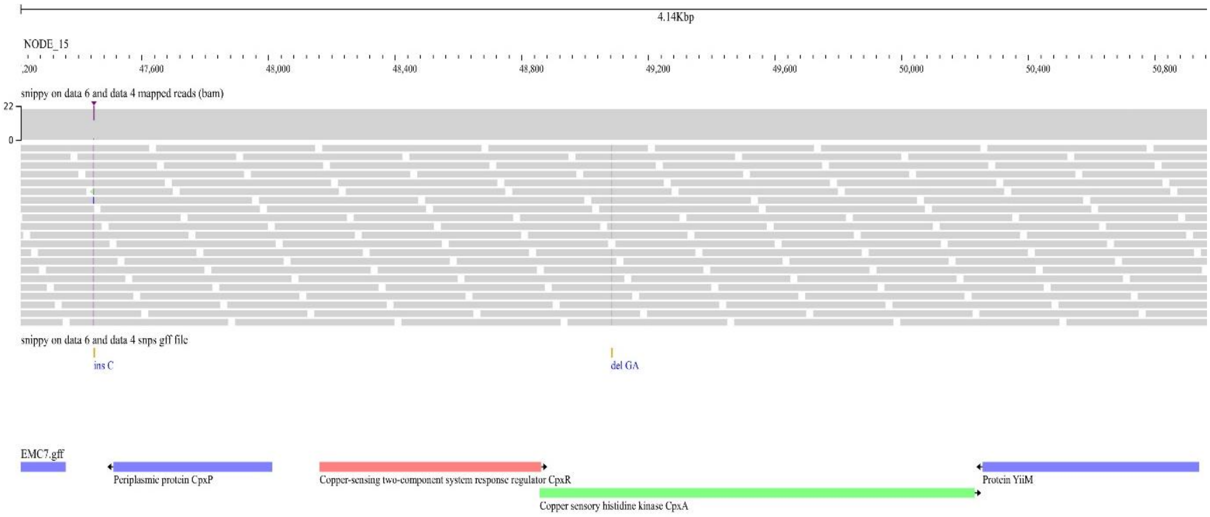


**Supplementary Figure S6a:** Multiple mutations (del GA, ins C) in *cpx* locus (*cpxP* and *cpxA*) in LTM01.

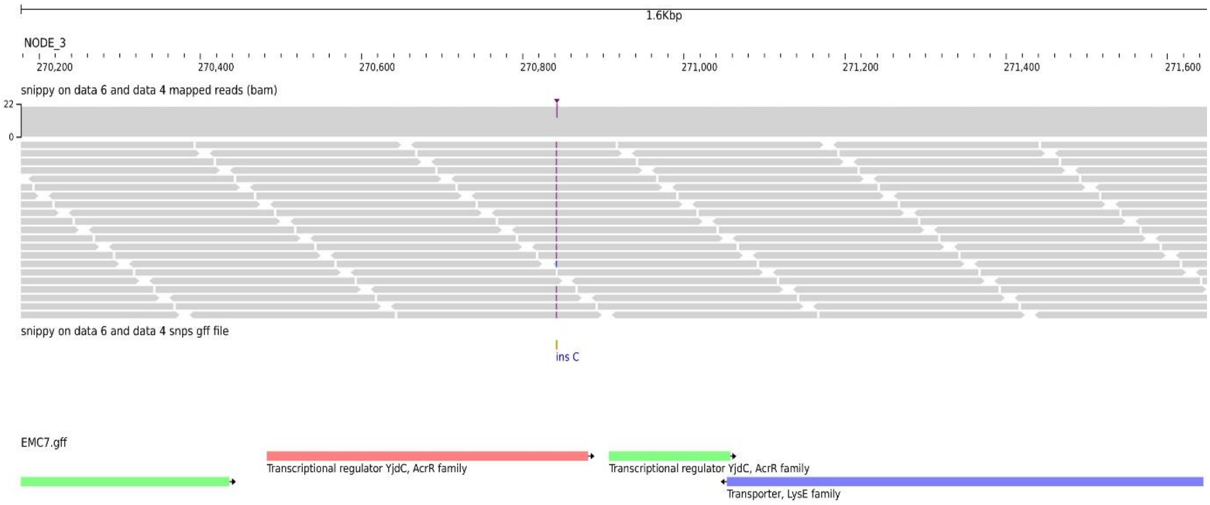


**Supplementary Figure S6b:** Ins C mutation in Transcriptional regulator gene *yjdC, acrR* family in LTM01.

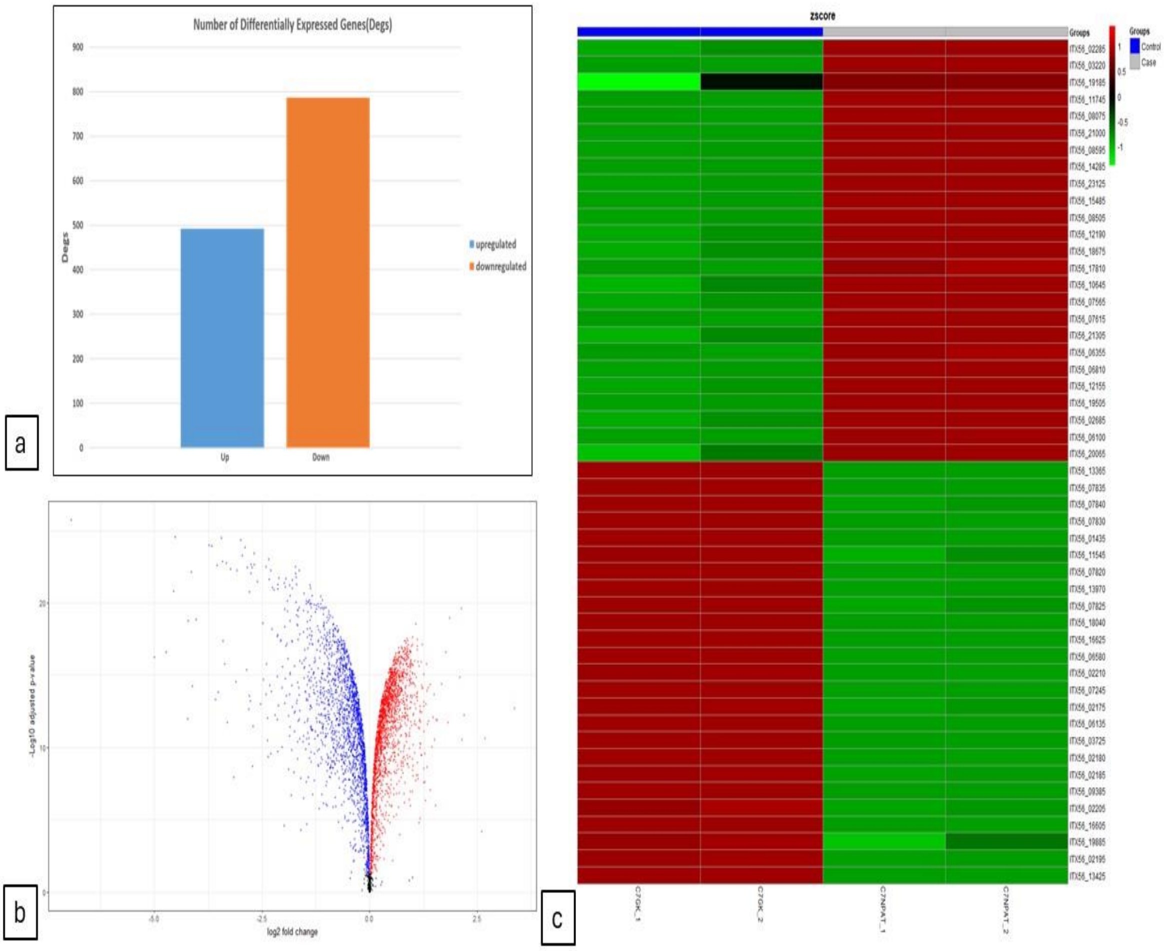


**Supplementary figure S7:** Differential transcriptomic response of wild-type *Leclercia barmai* EMC7 under glucose-3NPA (C7NPAT) relative to glucose-$KNO_3$ (C7GK).

(a) Bar plot showing the total number of differentially expressed genes identified in EMC7 under glucose-3NPA. A total of 1,279 genes were differentially expressed, including 492 upregulated and 787 downregulated genes.

(b) Volcano plot showing the distribution of differentially expressed genes based on log2 fold change and adjusted p-value. Red points indicate significantly upregulated genes, blue points indicate significantly downregulated genes, and grey points indicate genes not passing the applied differential-expression threshold. The plot shows broad transcriptional remodeling in EMC7 under glucose-3NPA, with a larger downregulated fraction than upregulated fraction.
(c) Heatmap showing z-score-normalized expression patterns of representative differentially expressed genes across glucose-$KNO_3$ control and glucose-3NPA treatment samples. The two glucose-$KNO_3$ samples clustered separately from the two glucose-3NPA samples, indicating condition-specific transcriptional restructuring under 3-NPA exposure.
